# Highly parameterized polygenic scores tend to overfit to population stratification via random effects

**DOI:** 10.1101/2024.01.27.577589

**Authors:** Alan J. Aw, Jeremy McRae, Elior Rahmani, Yun S. Song

## Abstract

Polygenic scores (PGSs), increasingly used in clinical settings, frequently include many genetic variants, with performance typically peaking at thousands of variants. Such highly parameterized PGSs often include variants that do not pass a genome-wide significance threshold. We propose a mathematical perspective that renders the effects of many of these nonsignificant variants random rather than causal, with the randomness capturing population structure. We devise methods to assess variant effect randomness and population stratification bias. Applying these methods to 141 traits from the UK Biobank, we find that, for many PGSs, the effects of non-significant variants are considerably random, with the extent of randomness associated with the degree of overfitting to population structure of the discovery cohort. Our findings explain why highly parameterized PGSs simultaneously have superior cohort-specific performance and limited generalizability, suggesting the critical need for variant randomness tests in PGS evaluation. Supporting code and a dashboard are available at https://github.com/songlab-cal/StratPGS.

## 1 Introduction

In recent years, polygenic scores (PGSs), defined as the sum of effects across a large number of genetic variants, have garnered wide interest and discussion regarding their application in research and clinical settings. The focus on PGSs stems from their success and availability. First, by amalgamating the effects of thousands, and potentially up to millions, of variants, some PGSs have performed as well as risk predictors arising from rare monogenic mutations, as measured by odds ratio and correlation between PGS and phenotype (1). Second, the convenience of being interpreted as a simple high-level disease predictor has led to the wide-spread use of PGSs as genetic risk instruments in consumer, research and clinical genetics settings (2; 3; 4). Third, in response to the growing literature on PGS modeling and data-analytic methodology for more generalizable PGSs (5), efforts have been made to curate PGSs alongside training methodology and metadata (6).

Despite their utility, PGSs have well-recognized issues, however. An overwhelming majority of PGSs are trained on cohorts of European ancestry; in particular, a recent study reported a 4.6-fold overrepresentation of individuals of European descent in PGSs compared to their global demographic proportion (7). Not only do PGSs trained on specific ancestries port poorly to other ancestries (8), but even PGSs trained for a cohort of the same ancestry may vary in performance across different subgroups stratified by age, sex or socio-economic status (9). There is also a lack of efficacious PGS methodology for individuals of multiple ancestries (i.e., admixed individuals), and the accuracy of a PGS may vary across the genetic ancestry continuum (10).

The poor cross-ancestry portability of PGSs tailored for specific ancestries has been attributed to inter-population differences in allele frequencies and linkage disequilibrium (LD) patterns (11; 12; 13; 14), closely related population-specific factors including genetic drift (15), and differences in environmental and social factors including social determinants of health (5). In contrast, variable cross-group portability of PGSs within the same ancestry has been linked to environmental stratification driven by demographic history (16) and confounding by non-genetic factors or assortative mating (17).

Another viewpoint may be obtained by considering why PGSs typically include large numbers of variants. Earlier studies have reported that, for typical traits, the most significantly associated loci explain only a modest fraction of the estimated genetic variation, and that including more variants with non-significant effects would help explain “missing heritability” (18; 19). Together with evidence that complex trait variation is only weakly explained by core genes alone — which presumably cluster by disease-relevant pathways — a perspective has emerged that the genetic etiology of many complex traits is characterized by a large number of variants that individually contribute small effects (20; 21). This is consistent with the successes of PGSs that include non-significant variants (highly parameterized PGSs) across biomedical phenotypes such as neurodegenerative disorders (22), cardiovascular traits (23; 24; 25; 26) and respiratory diseases (23).

Here, we present a new mathematically motivated perspective on PGSs, which challenges the notion that non-significant variants contributing to PGS predictability are causal yet non-significant at a genome-wide level owing to small effects. We argue that highly parameterized PGSs are often over-parameterized, with the over-parameterization capturing genetic structure of the discovery cohort (i.e., population stratification), which in turn, may boost performance within the same population as the discovery cohort but not generalize well to other populations. Using mathematical models and empirical validation across 141 quantitative phenotypes from the UK Biobank (UKB), we demonstrate that, at least for the most part, non-significant variants included in PGSs do not capture causal signals but rather population structure. Our results carry implications for principles underlying generalizable PGSs for science and health equity, motivate new tests for stratification-driven overfitting of PGSs, and provide a justification for ad-hoc overfitting diagnostics that have appeared in the literature.

## 2 Results

### Over-parameterization in highly parameterized PGSs: mean corpuscular haemoglobin as a test case

Highly parameterized PGSs tend to improve performance by incorporating variants that are not genome-wide significant (Figure 1A), allegedly owing to the inclusion of small-effect causal variants. Consequently, PGSs are typically constructed using lenient variant inclusion thresholds by design. In fact, across more than 3, 600 PGSs reported in the PGS Catalogue (6), the median number of variants included in a score is 5, 578 (Figure 1B), with a maximum number of variants reported exceeding 10^7^. This is more than a 100-fold greater than the average number of GWAS hits across phenotypes, which is estimated to be between 10 and 50 (27; 28).

**Figure 1:**
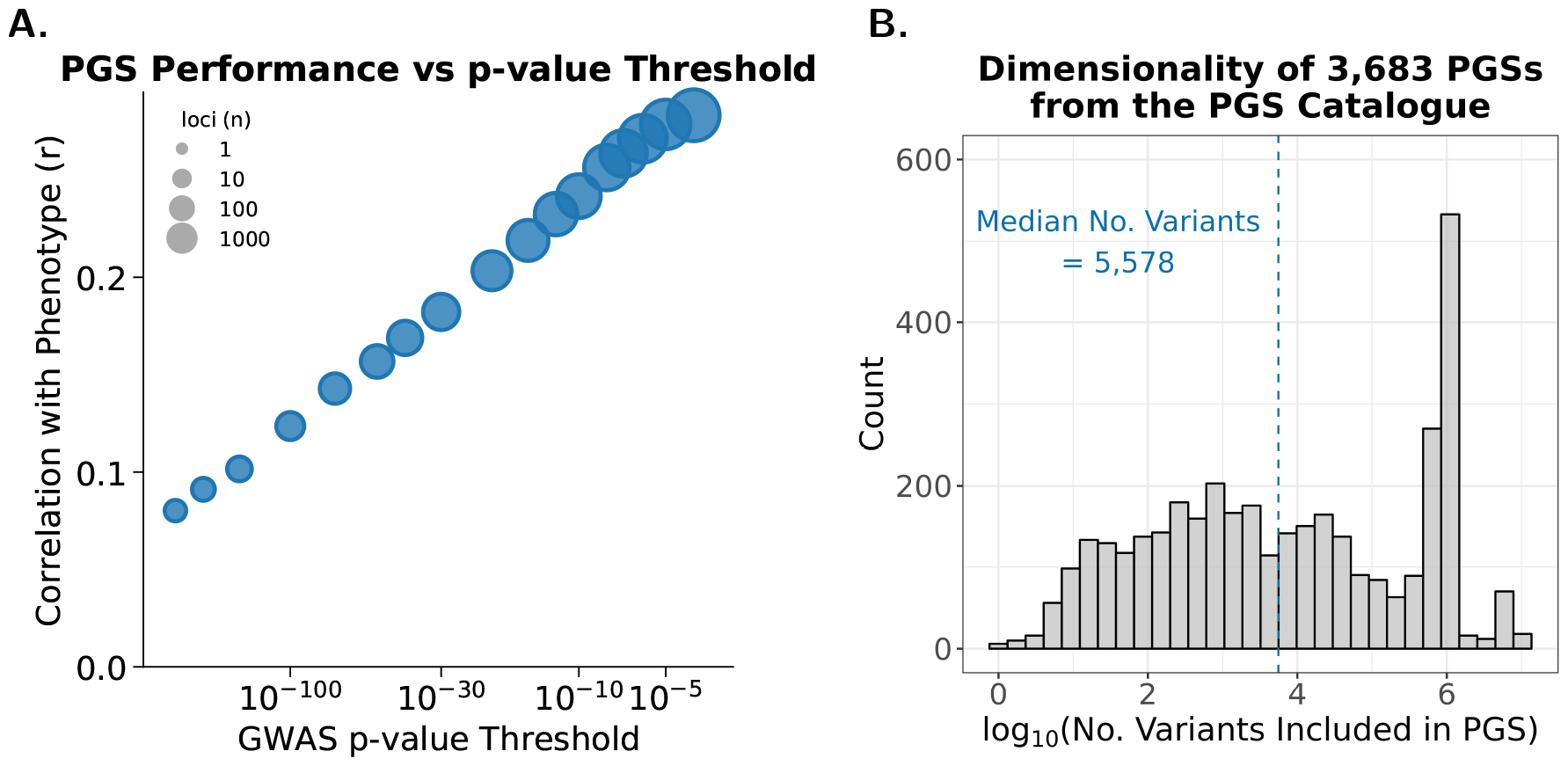
**A**. Improvement in PGS performance with increasing *p*-value threshold used for including variants in the clumping and thresholding PGS algorithm (1). Performance measured by mean Pearson correlation with phenotype across up to 141 quantitative traits in a test cohort of *n*_test_ = 68, 931 individuals from the UKB (Materials and Methods). **B**. Histogram of log-counts of the number of variants included in PGS for all 3, 683 PGSs reported in the PGS Catalogue with available metadata.

However, the design of highly parameterized PGSs seems statistically unsound, as we would expect the vast majority of non-significant variants under a lenient *p*-value threshold to be neither causal nor systematically associated with the phenotype, especially when using large cohorts with hundreds of thousands of individuals. We consider this arguably more conservative statistical view, which renders the effects of genome-wide non-significant (hereafter “non-significant”) variants merely random, as one would get from any statistic under the null distribution.

In order to motivate our perspective with a concrete example, we first consider 15 PGSs for mean corpuscular haemoglobin (MCH), a biomarker for anaemia, and evaluate them using data from the UKB (Supplementary Material S5). These PGSs were obtained from the PGS Catalogue (6), and they differ in the number of variants included, study cohorts and training methodology (29; 30; 24; 25; 26) (Supplementary Table S4). Using a set of 11 performance metrics of PGS (Supplementary Table S1), we compare the relative performance of each original MCH PGS against 100 permuted PGSs (pPGSs), which we obtained by randomly permuting the effects among non-significant variants included in the original PGS (Materials and Methods). These variants have distinct effects, so permuting them changes the PGS. Importantly, if the non-significant variants reflect real biological effects, then the PGSs are expected to be *sensitive* to permutations and pPGSs should therefore demonstrate deterioration in performance. Conversely, if the non-significant effects are indeed purely random, one would expect the PGSs to be *insensitive* to permutation and the performance of a pPGS to be on par with its corresponding original PGS.

Of the 15 MCH PGSs (Supplementary Material S5), only 9 report sensitivity to permutation (Supplementary Figure S4). That is, for the remaining 6 PGSs, the net effect of non-significant variants is not generally better than random effects via permutation, which is consistent with the conservative statistical view and implies over-parameterization of these PGSs. Yet, we observe stronger performance for MCH PGSs that include larger numbers of variants (Supplementary Figure S3), which is in line with the same observation across various phenotypes (Figure 1A). Intuitively, it is contradictory to observe improved performance for highly parameterized PGSs, and at the same time, insensitivity of these PGSs to permutation, which implies redundancy of variants. In particular, the best performing PGS is the one including the most variants among all 15 MCH PGSs, yet, it is also among the 3 most insensitive PGSs, indicating over-parameterization (Supplementary Figure S4A; Supplementary Material S5). A natural question is to consider how over-parameterization due to the inclusion of random effects coincides with improved PGS performance.

### Mathematical theory justifies the statistical perspective

For a polygenic effect vector ***β*** = (*β*_1_, …, *β*_*p*_) for some trait and an individual-by-genotype matrix **X** with the same *p* variants as in ***β***, we consider **X*β*** as the PGS. It can be shown (Materials and Methods) that, under simplifying assumptions, the following relationship holds up to a scaling factor for any ***β***:

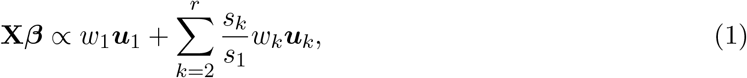

where *r* = min{*n, p*}; ***u***_*k*_ and *s*_*k*_ respectively denote the *k*-th left singular vector and corresponding singular value of **X** (with *s*_1_ being the largest); and *w*_*k*_ is a scalar weight.

The singular vectors and singular values of the genotype matrix in Eq. (1) are fixed and independent of the polygenic effect vector ***β***, which determines the weights *w*_*k*_. Concretely, *w*_*k*_ is proportional to the cosine similarity between ***β*** and the *k*-th principal direction of **X** (Materials and Methods). Therefore, in the extreme case where the effects in ***β*** are all random, *w*_1_, …, *w*_*r*_ are expected to follow the same null distribution, and **X*β*** is expected to be primarily determined by the fixed ratios *s*_*k*_*/s*_1_. The top singular values from genetic data capture more variation than subsequent axes (31; 32). In the case of random effects ***β***, the PGS **X*β*** is thus expected to be correlated with the top principal components (PCs) of the genotype matrix. Consequently, for traits correlated with population structure, random PGSs are expected to capture the trait better than expected by random chance alone.

In practice, some of the non-significant effects in a PGS may not reflect random values but rather real variant effects. Yet, we can restrict Eq. (1) to the random part of the PGS and the rationale above would still hold. This mathematically explains why we expect that highly parameterized PGS models tend to improve performance, as observed in the MCH example. The extent to which this phenomenon applies to a specific trait is expected to depend on the correlation of the trait with the population structure of the fitted cohort. Eq. (1) also suggests that a stronger correlation of the PGS with the population structure tagged by the *k*-th principal direction of the data will lead to a higher weight *w*_*k*_. This observation suggests that the empirical distribution of *w*_1_, …, *w*_*r*_ can be informative of population stratification biases in a PGS.

Our perspective offers a mathematical explanation for the enhanced performance observed for highly parameterized PGSs. This aligns with a conservative statistical viewpoint, suggesting that most non-significant variants in PGSs are unlikely to consistently influence trait variation across populations or correspond to a stable or generalizable molecular mechanism. Importantly, our explanation also corroborates the poor portability observed in PGSs. Particularly, the inclusion of random effects in a PGS may lead to similar performance across populations that present similar correlation structure between the trait and population stratification. In contrast, when transferring a PGS between populations that present different correlation structures with the trait, the random effects are expected to contribute differently to performance. We next evaluate our perspective empirically, by considering various kinds of PGSs where some or all of the effects are random, which we collectively refer to as *random projections* (Figure 2; Materials and Methods). Our extensive study of these random projections reveals how prevalent our mathematical observations are when considering a large set of traits.

**Figure 2:**
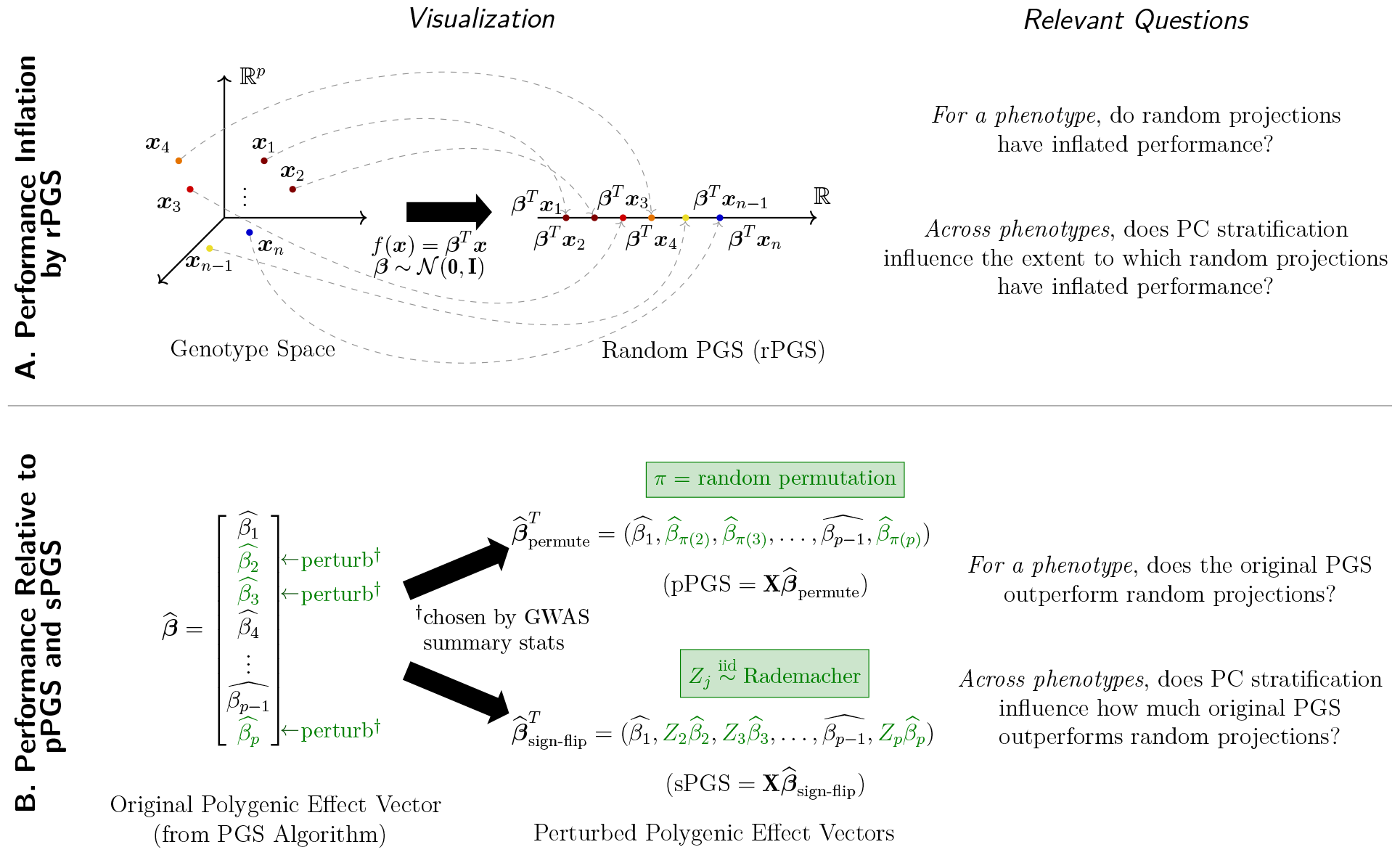
Summary of random projection experiments performed on UK Biobank (UKB) data. **A**. Performance Inflation by rPGS. **B**. Performance Relative to pPGS and sPGS. Perturbed variants (*†*) are determined by whether the GWAS *p*-value with the phenotype exceeds a user-specific threshold, *p*_non-sig_. We consider *p*_non-sig_ ∈ {10^−6^, 10^−7^, 10^−8^, 10^−10^} in our work.

### Random PGSs can explain phenotypic variation across a wide range of traits

We begin by considering a set of 141 quantitative traits measured for *n* = 487, 296 individuals in the UKB (Materials and Methods). To evaluate whether random effects can be used to explain some of the variation of these traits, we first generated 200 random PGSs (rPGSs) using effect sizes we sampled from a standard normal distribution for a randomly selected 10% of the variants in the data (Figure 2A; Materials and Methods). The rPGSs were not directly tied to any of the traits and we evaluated the extent to which each rPGS captured the variation of each trait. Specifically, for a given trait, we evaluated the performance of each rPGS by measuring the change in *R*^2^ when adding the rPGS to a baseline linear regression model that factored in age, sex, and the top 20 PCs of the genotype data. For each of the 141 traits, we computed mean incremental *R*^2^ values for the performance of the rPGSs.

Next, we evaluated whether the performance of the rPGSs is better than what could be expected by chance. We permuted each trait 100 times across individuals and repeated the above procedure for calculating mean incremental *R*^2^ for each permuted trait based on the same set of 200 rPGSs. This allowed us to compare the performance of the rPGSs for each original trait against its null distribution and calculate empirical *p*-values. We found 126 (89%) traits with empirical *p*-value *<* 0.05 (Figure 3A). Repeating our analysis on inverse rank-normalized (IRNT) versions of the 141 traits revealed qualitatively similar results, with 135 (96%) traits with empirical *p*-value *<* 0.05.

**Figure 3:**
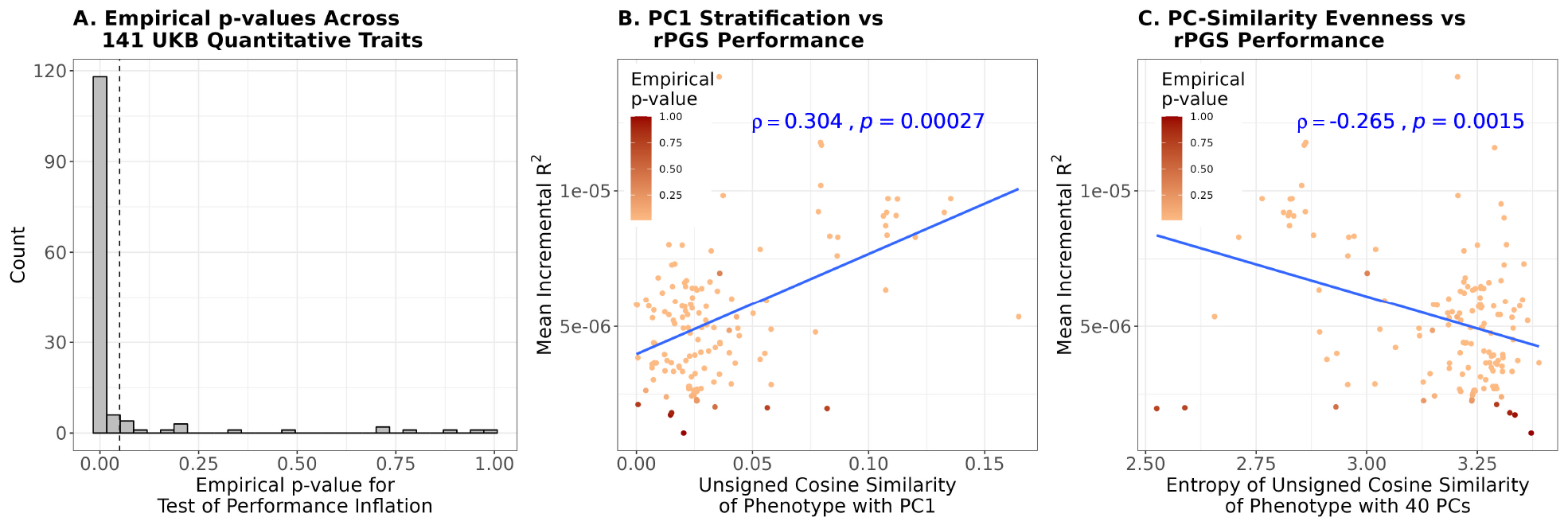
**A**. Distribution of empirical *p*-values under the null hypothesis that random PGSs do not explain phenotypic variation. **B**. Relationship between mean incremental *R*^2^, across rPGSs, for a phenotype against its unsigned cosine similarity with PC1. **C**. Relationship between mean incremental *R*^2^, across rPGSs, for a phenotype against the entropy of unsigned cosine similarities with the top 40 PCs, a measure of evenness of the relationship between the phenotype and the PCs. PGS performances are evaluated with respect to original phenotypes. Full results for both original and inverse rank-normalized versions are summarized in Supplementary Figure S5.

Our mathematical analysis suggests population stratification as the source of the inflated performance of rPGSs. Population structure captured by rPGS can only inflate performance if the trait itself is also correlated with population structure. Moreover, the level of inflation of performance is expected to depend on the extent to which the trait is correlated with population structure. In order to quantify the latter, we considered two measures of population stratification of a phenotype. First, motivated by our results from Eq. (1), we considered the (unsigned) cosine similarity between the phenotype and the first PC of the genotypes as a measure of how much a trait is correlated with population structure. Indeed, we observe a significant positive rank correlation between the latter and the incremental *R*^2^ (Spearman *ρ* = 0.3, *p* = 2.7 *×* 10^−4^; Figure 3B). As a second evaluation metric, we considered the entropy of unsigned cosine similarity across the first 40 PCs of the genotypes, which quantifies divergence from a uniform concordance between the phenotype and the PCs (as expected in the absence of correlation of the phenotype with population structure). As expected, we observed a significant negative correlation between the latter and incremental *R*^2^ (Spearman *ρ* = −0.26, *p* = 1.5 *×* 10^−3^; Figure 3C). Altogether, our results link the performance of random effects to the correlation between the phenotype and population stratification.

### PGSs constructed using liberal variant inclusion thresholds overfit to population structure

Our initial observation with MCH showed that including non-significant variants in a PGS is generally not better than assigning random effects to such variants. We next explored how widespread this phenomenon is across a large number of traits. We have just seen that information captured by random effects is driven by the phenotype’s stratification by PCs. We assessed how the out-of-sample performance of a trained PGS with non-significant variants might be influenced by overfitting to population stratification introduced during training.

We trained and evaluated PGS models for a subset of 103 quantitative traits based on individuals of (self-identified) European descent from the UKB data, where the subset of traits was chosen based on sufficiently many variants with non-significant GWAS *p*-values (Materials and Methods). GWAS summary statistics were obtained using age, sex and the top ten genetic principal components as additional covariates to control for possible confounding. For each trait, we applied clumping and thresholding (C&T) on the GWAS summary statistics (Materials and Methods), and reweighed effect sizes using multiple regression. We obtained a *lenient* model using variants with *p*-value *< p*PGS = 10^−5^ and a training set of *n* = 288, 728 individuals. We applied this model to predict PGS for a held-out test set of *n* = 68, 931 individuals. As in the MCH analysis, we evaluated the performance of the PGS of each trait by considering 100 pPGSs we computed by permuting the effects among non-significant variants (those with *p*-value ∈ [*p*_non-sig_, *p*PGS), where *p*_non-sig_ is a user-designated perturbation cutoff). In addition, we considered 100 sign-flipped PGSs (sPGSs), which we obtained by randomly flipping the directionality of non-significant variants (Figure 2B). A PGS with non-significant variants that are more informative than random effects — such as the permuted and sign-flipped effects just described — is expected to tend to perform better than pPGSs and sPGSs and thus report low empirical *p*-values.

Considering variants whose GWAS *p*-values lie in the range [10^−8^, 10^−5^) as non-significant (i.e., *p*_non-sig_ = 10^−8^), we evaluated for each PGS whether it performs better than the same PGS only with random effects in place of non-significant effects (i.e., pPGSs and sPGSs). Measuring performance using percentile-prevalence rank correlation, we found that only 29 (resp., 25) of the 103 PGSs show statistical evidence (empirical *p*-value *<* 0.05) of improvement over pPGSs (resp., sPGSs) (Figure 4A). Constructing pPGS and sPGS with even weaker variants led to even fewer PGSs showing improvement over PGSs with random effects. In particular, for *p*_non-sig_ = 10^−6^, we found the number of PGSs with empirical *p*-value *<* 0.05 to be no different from that expected by chance (Figure 4A). Consistent with our mathematical perspective, these results show that the inclusion of low significance variants in a PGS generally does not improve performance over random effects induced by permutation or sign flipping of effects.

**Figure 4:**
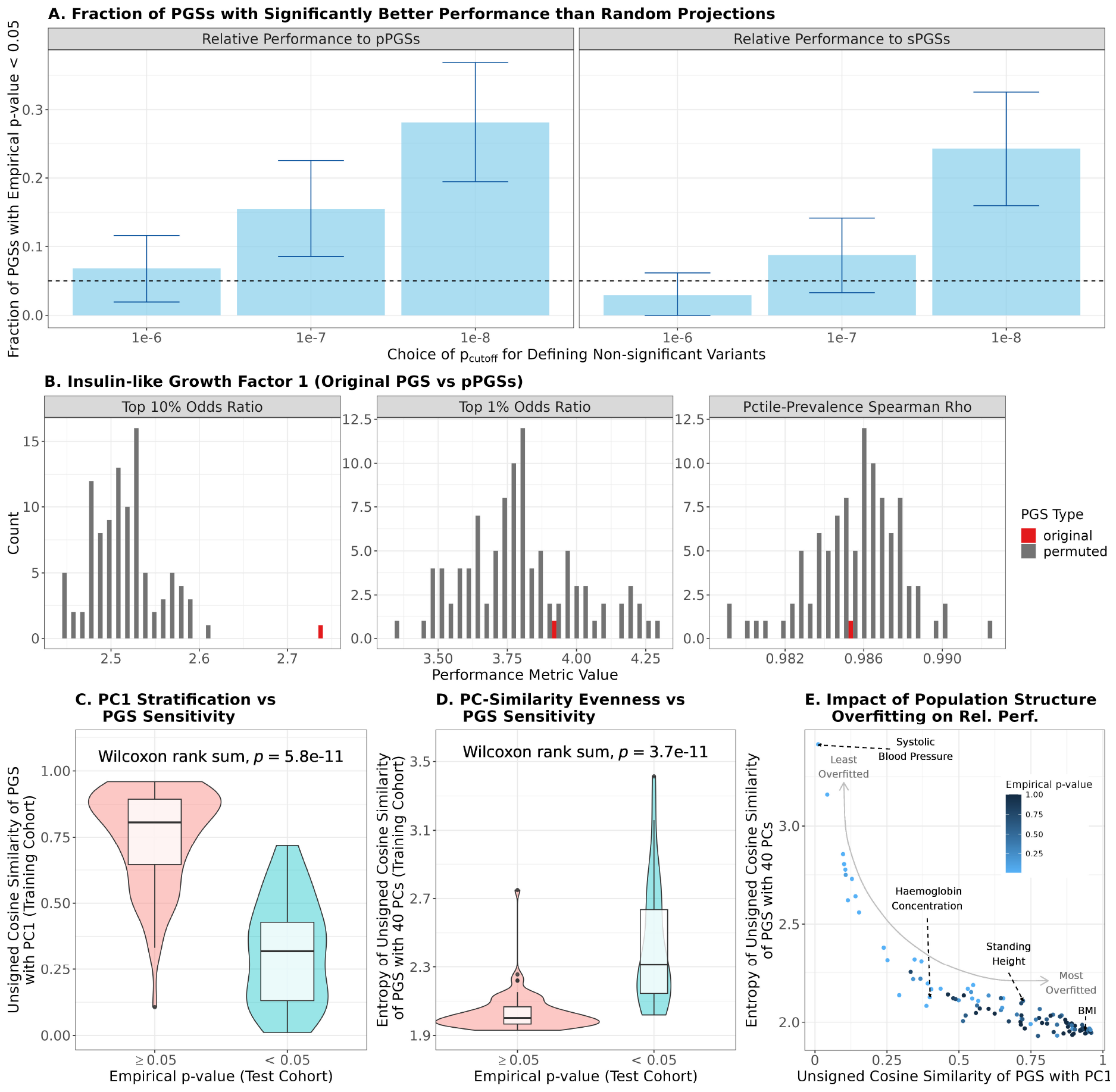
Performance of PGSs relative to random projections. **A**. Fraction of PGSs with relative performance test empirical *p*-value *<* 0.05, where performance is measured by percentile-prevalence rank correlation and various choices of *p*_non-sig_ are used to determine non-significant variants (i.e., variants with GWAS *p*-value at least *p*_non-sig_). Approximate 95% confidence intervals are computed from the empirical fraction, assuming uniform Type I Error across phenotypes. **B**. Empirical histograms of performance computed for Insulin-like Growth Factor 1 original PGS and pPGSs (non-significant variants determined by *p*_non-sig_ = 10^−8^), across multiple metrics, on the test cohort. **C**. Distribution of cosine similarities of each phenotype’s original PGS with PC1 in the training cohort. Distributions are split by empirical *p*-value (relative performance to pPGSs with *p*_non-sig_ = 10^−8^), and performance is measured by cosine similarity with phenotype. **D**. Distribution of entropies of cosine similarities of each phenotype’s original PGS with PC1 in the training cohort. Distributions are split by empirical *p*-value, and performance is measured by cosine similarity with phenotype. **E**. Scatterplot of entropies of cosine similarities and cosine similarities with PC1, computed for all 103 PGSs. Points coloured by empirical *p*-value, with performance measured by cosine similarity with phenotype.

In order to evaluate the contribution of random effects in PGS models with a more stringent variant inclusion criterion, we further evaluated a set of 68 traits for which at least five variants with GWAS *p*-value ≥ 10^−10^ exist. This allowed us to ask whether not just the lenient models, but even more stringent ones can have their least informative variants showing no improvement over random effects and thus be insensitive to pPGS and sPGS perturbations. Indeed, considering only variants with *p*-value *< p*PGS = 10^−8^ in the PGS models (instead of *p*PGS = 10^−5^ for lenient models) and setting the non-significance cutoff to *p*_non-sig_ = 10^−10^ for pPGS and sPGS construction, we found no evidence that the original PGSs generally improves performance over models containing random effects (Supplementary Figure S6).

Re-evaluating the sensitivity of the PGSs to pPGSs and sPGSs under different PGS performance metrics presented great variability in sensitivity for both the lenient and the more stringent PGSs (Table 1; see Figure 4B for example of PGS for Insulin-like Growth Factor 1). Interestingly, we found that correlation metrics demonstrate very high sensitivity to pPGS and sPGS perturbations, indicating that random effects are not as predictive as the non-significant effects in the original PGS models. This is potentially owing to overfitting introduced during the effect size reweighting process of fitting the PGS models.

**Table 1:** Numbers of PGSs with empirical *p*-value *<* 0.05 (i.e., significantly more performant) based on different non-significance *p*-value thresholds *p*_non-sig_ for constructing pPGS, across various PGS performance metrics. Original PGSs were built from C&T using a lenient GWAS threshold of *p*_PGS_ = 10^−5^ for inclusion of variants. For the rightmost column, PGSs were built from C&T using a stringent threshold of *p*_PGS_ = 10^−8^, and each entry reports the number of PGSs out of 68 considered phenotypes.

| Metric | Lenient Model<br>Out of 103 Phenotypes |  |  | Stringent Model<br>Out of 68 Phenotypes |
| --- | --- | --- | --- | --- |
| | $p_{\text{non-sig}} = 10^{-6}$ | $p_{\text{non-sig}} = 10^{-7}$ | $p_{\text{non-sig}} = 10^{-8}$ | $p_{\text{non-sig}} = 10^{-10}$ |
| Cor($y$, PGS) | 96 | 102 | 103 | 62 |
| $\rho(y, \text{PGS})$ | 103 | 102 | 103 | 63 |
| CosineSim($y$, PGS) | 8 | 18 | 26 | 8 |
| Top 10% Prevalence | 16 | 50 | 77 | 15 |
| Top 1% Prevalence | 3 | 11 | 24 | 2 |
| Top 10% Odds Ratio | 16 | 49 | 78 | 15 |
| Top 1% Odds Ratio | 3 | 11 | 24 | 2 |
| Cor(Percentile, Prevalence) | 5 | 21 | 38 | 4 |
| $\rho(\text{Percentile}, \text{Prevalence})$ | 7 | 16 | 29 | 5 |
| Cor(Percentile, Ave. $y$) | 7 | 13 | 29 | 5 |
| $\rho(\text{Percentile}, \text{Ave. } y)$ | 4 | 12 | 35 | 7 |

Lastly, to quantify the extent to which a PGS is overfitted to population structure during training, we computed 12 additional measures of population stratification of a PGS (Materials and Methods). These include the unsigned cosine similarity between the PGS and the first PC of the genotypes as well as the entropy of unsigned cosine similarity across the first 40 PCs of the genotypes, calculated on the training cohort. Across 103 lenient PGSs with *p*_non-sig_ = 10^−8^, PGSs with empirical *p*-values *<* 0.05 (reflecting significantly better performance of the original PGS relative to pPGSs, where PGS performance is measured using PGS-phenotype cosine similarity) and those with empirical *p*-values ≥ 0.05 exhibit a significant difference in how well the PGS is correlated with the first PC (Figure 4C). For the same performance metric, we also observe a significant depletion in the entropy of unsigned cosine similarity of non-significant empirical *p*values relative to significant ones (Figure 4D). These statistical relationships hold regardless of the choice of *p*_non-sig_ (Supplementary Figure S7); whether sPGSs or pPGSs is computed (Supplementary Figure S7); and whether stringent PGSs (with *p*PGS = 10^−8^) are considered instead (Supplementary Figure S9). Taken together, these results demonstrate that overfitting PGSs to population structure during training explains their low out-of-sample performance relative to PGSs with random effects assigned to non-significant variants (Figure 4E).

### Additional analyses of PGS insensitivity

Delving deeper, we next investigated whether PGSs were generally more sensitive to pPGSs or to sPGSs (Supplementary Material S6). Broadly, we observed more significant empirical *p*-values when evaluating the performance of original PGSs relative to pPGSs than to sPGSs (Supplementary Figures S11 and S12). However, the strength of this relationship also depends on the choice of performance metric.

We also investigated if patterns related to either the set of non-significant variant effects or the complementary set of fixed variants were predictive of PGS insensitivity (Supplementary Material S7). Significant positive rank correlations were observed, across PGSs, between the sum of absolute value of effect sizes of non-significant variants and the empirical *p*-values of the original PGS against random PGSs. For example, measuring performance using Top 10% odds ratios yielded Spearman *ρ* ∈ [0.37, 0.72]. This is regardless of how non-significant variants were called, whether a lenient or stringent PGS was considered, or whether performance was relative to pPGS or sPGS (see Supplementary Figures S13 and S15). On the other hand, no significant relationships were detected between the empirical variance of effects of non-significant variants and empirical *p*-values. These findings show that the performance of the original PGS relative to pPGSs or sPGSs is not driven by variation in the non-significant effects, as one might posit as an *a priori* statistical explanation. It is instead driven by the magnitude of effects of non-significant variants, suggesting that the larger the (random) non-significant effects are, the greater they might account for overfitting to population stratification and reducing out-of-sample performance.

## 3 Discussion

Our present work investigates the impact of population stratification on PGS performance inflation under the view that PGSs are summations of variant effects, a considerable number of which may be random. Combining mathematical reasoning with empirical validation on the UKB, we have found that highly parameterized PGSs capture population structure, as proxied by PCs. Such a phenomenon leads to inflated performance by random projections constructed via assigning random effects to variants (rPGSs), as well as low sensitivity of trained PGSs to perturbations of their constituent variants’ effects (pPGSs and sPGSs). Notably, our analysis of MCH PGSs suggests that up to 40% of currently reported trait-specific PGSs may face such an over-parameterization issue. These results challenge the view that the strong performance of highly parameterized PGSs is principally explained by numerous non-significant variants contributing small but causal marginal effects.

The reasoning behind random projections capturing population structure can be traced back to earlier work in high-dimensional statistics, specifically in the use of random projections to reduce data dimensions while preserving pairwise distances within the sample (33). While the earlier results do not directly translate to our setting, the fact that the individuals comprising a cohort share genetic distances that proxy ancestral or genealogical divergences and that the distances can be approximated in a lower-dimensional space *via the use of statistical noise*, is appreciable. Our present work has demonstrated that such a phenomenon applies to biobank-scale data.

Our framework is similar to, and justifies, previously reported tools for diagnosing population stratification of PGSs, which are scarcely used beyond a few selected works. These include a downstream diagnostic computing rank correlations between the polygenic effect vector and the SNP loadings vector of each PC (34), and an upstream diagnostic computing random PGSs using approximately independent GWAS-insignificant variants (i.e., *p*-value *>* 0.5) (35; 36). Our framework differs from these tools in several technical ways (for example, how and why we choose variants to construct random projections), and also augments the toolkit for diagnosing population stratification-related biases (for example, the use of random permutations and sign flips to assess relative performance).

Our work has a few limitations. First, in our analysis of MCH PGSs, the overlap between our evaluation cohort and each score’s training cohort was not explicitly considered. A more fair approach would be to analyze the sensitivity of these PGSs with respect to a cohort of individuals external to the combined set of individuals making up each training cohort. However, the fact that a considerable fraction of the PGSs appear insensitive despite cohort overlap suggests that such a comparison may reveal even more insensitive PGSs. Second, our observation that PGS insensitivity to and predictability inflation by random projections are both driven by population stratification is contingent on the use of C&T as a PGS construction algorithm. It is possible that other PGS algorithms incorporating flexible modeling strategies will sufficiently capture or correct for population stratification, resulting in the attenuation or absence of the trend. It is also possible that the inclusion of more PCs as covariates when performing GWAS (i.e., greater than 10, which we have used in our work) will sufficiently correct for population stratification. An extensive investigation into other PGS algorithms and PC correction strategies would clarify this matter.

Our findings carry important implications for interpreting and evaluating PGSs. First, our finding that non-significant variants capture population structure may appear to contradict the widely-held view that many complex traits carry large numbers of variants each contributing a small marginal effect. This latter view may still be true, although our analyses suggest that even biobank-scale cohorts are insufficiently powered to estimate variant effects to a high precision, with the inferred effects capturing population structure of the discovery cohort instead. PGSs that are derived from flexible probabilistic models may overcome this issue by inferring more accurate effect distributions for these variants, as evidenced by our observation that some MCH PGSs are sensitive to random projections. However, it remains unclear what fraction of publicly available PGSs suffer from over-parameterization capturing population stratification. A recent study on the UKB reported that PGSs can be significantly correlated with geography despite adjustments for population structure, reinforcing the difficulty in distinguishing between molecular-level and population-stratification explanations of the strong prediction accuracies observed of PGSs (37). In any case, we believe that the performance inflation and relative performance audits presented in this work are valuable additions to standard performance metrics, to ensure that the effects of non-significant variants in a PGS are reliably estimated or tuned post-GWAS.

Second, through computing various metrics to measure population stratification, PGS performance and summary statistics related to the constituent variants of a polygenic effect vector (i.e., Perturbed-Fixed Architecture in Table 2 of Materials and Methods), we have identified some metrics that appear more useful than others at detecting PGS insensitivities. The greater incidence of PGSs with low sensitivity to percentile-related metrics is consistent with recent work demonstrating the large uncertainty in individual PGS estimation, which would naturally impact metrics relying on groupings of individuals by risk strata (38). Our finding of low cosine similarity of a PGS with phenotype, relative to random projections, being well-predicted by PC stratification is particularly novel (Figure 4C-E), given that the use of cosine similarity as a PGS performance metric appears scarce to our knowledge. In general, practitioners may improve the efficacy of sensitivity analyses by computing the more discerning metrics identified in our work. In our application to MCH PGSs, we have also observed that no one metric consistently selects the “most performant” PGS. While this may be particular to the trait we have studied, our observation entails the potential utility of investigating the use of a diverse set of metrics simultaneously in evaluating PGSs.

**Table 2:** Key concepts used throughout our work.

| Concept | Definition | Examples |
| --- | --- | --- |
| Random Projection | linear combination of allelic dosages, with some / all weights drawn randomly | $x_1\beta_1 + \dots + x_p\beta_p, \beta_j \stackrel{\text{iid}}{\sim} N(0, 1)$<br>(see also Figure 2) |
| Performance<br>(of PGS) | how well a PGS predicts a phenotype | correlation between PGS and phenotype,<br>percentile-prevalence correlation |
| PC Stratification<br>(of PGS or phenotype) | how strongly a PGS or a phenotype is stratified by the principal components of a cohort of interest | correlation between PGS and PC1, cosine similarity of PGS and PC1 |
| Perturbed-Fixed Architecture<br>(of PGS) | quantities calculated from effects of either the set of (GWAS) non-significant variants or the set of significant variants in a PGS | empirical variance of effects of variants with GWAS $p$-value $>10^{-6}$ |

Given the implications of our findings, we note several actionable next steps to advance the operationalization of trustworthy PGSs. First, our random projection sensitivity checks can be applied to trait-specific PGSs other than MCH, which are trained using various methodologies and on different cohorts, to evaluate the extent to which highly parameterized polygenic effect vectors are capturing small but precise variant effects (which would support their veridicality) rather than population structure of the training cohort. Second, our sensitivity checks select variants whose *p*value exceeds a user-specific threshold. This approach may be limited by the availability of summary statistics for a desired cohort, especially given that many GWASs remain biased toward cohorts with European ancestry. In selecting variants to evaluate PGS sensitivity, one may also select variants based on external functional annotations, given the possibility that particular annotations are known to map biologically to a phenotype of interest. There is growing interest in variant effect predictors (VEPs) developed from advanced machine learning methods (39); such tools can be used to evaluate the precision of effects of classes of variants in a PGS defined by their predicted functional impact. More generally, sensitivity analyses can be used to interrogate the precision of the effects of variants at loci of interest.

Finally, our work contributes to ongoing discussions about improving personalized genomic predictions. Given the demonstrated utility of PGSs in identifying high-risk individuals across diseases, our work suggests that part of this success may be owing to the PGS capturing population structure, which implies the usefulness of population structure as a predictive signal. One promising direction is the development of methods that accurately disentangle ancestry-relevant and phenotype-relevant signals, thereby leveraging the utility of both to improve PGS prediction power (40). Ultimately, the inclusion of diverse individuals in GWAS and in training PGSs will improve personalized genomic predictions. However, given that efforts to collect data on more ancestrally diverse human populations have yet to mature, current PGSs should be audited to prevent overfitting to population structure, which would improve their generalizability. We believe that our random projection approach is an important step forward.

## Data availability

Results and scripts used for running our experiments are available at the Github repository https://github.com/songlab-cal/StratPGS.

## Acknowledgements

We thank Andy Dahl, Jonathan Flint and members of the labs of Iain Mathieson and Ziyue Gao for helpful discussions. The work described in this manuscript was approved by the UKB under application number 33751. This research is supported in part by an NIH grant R35-GM134922 and grant number CZF2019-002449 from the Chan Zuckerberg Initiative Foundation.

## 4 Materials and Methods

Our work rests on a few key unifying concepts that we define and summarize in Table 2. The first concept is a *random projection*, which encompasses random PGSs (rPGSs) as well as permuted PGSs (pPGSs) and sign-flipped PGSs (sPGSs). Next, *performance* refers to how well a PGS predicts or explains a phenotype. In this paper, we rely on up to 12 metrics to measure performance (Supplementary Table S1). Given a PGS or phenotype, its alignment with population structure is quantified by various measures of *PC stratification*, and we define and compute 12 such measures in our paper (Supplementary Table S2). Finally, for a polygenic effect vector ***β***, quantities measuring statistical patterns related to either the set of non-significant variants or the complementary set of fixed variants are computed, which we collectively term *perturbed-fixed architecture* (Supplementary Table S3).

### Mathematical Framework

Suppose a cohort consisting of *n* individuals genotyped across *p* variants is provided for PGS construction. Many PGS methods rely on two quantities dependent on the genotype matrix **X** ∈ R^*n×p*^: (1) principal components (PCs) computed from a subset of variants to estimate population-structural confounders; and (2) genome-wide association studies (GWAS) summary statistics obtained from association mapping to select variants for inclusion in the model. The latter quantities typically rely on the former PCs as covariates in the regression procedure, and also depend on the phenotype vector ***y*** measured across the individuals. To proceed with our mathematical treatment, we make two additional simplifying assumptions.

1. The set of *p* variants making up **X** are those used in constructing the PCs, and these variants moreover include all variants selected for inclusion during PGS training.
2. The PGS is computed on the normalized version of **X**. That is, **X** is obtained from the original genotype matrix of allelic dosages, by subtracting the *j*th column by 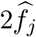 before scaling it down by 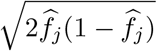 where 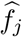 is the cohort’s (estimated) allele frequency.

Note that these assumptions are violated in practice, and we only require them for stating and deriving the key mathematical results that motivate our empirical study. (See Supplementary Material S2 for a discussion.)

Under our simplifying assumptions, the PGS vector can be expressed as a weighted sum of the PCs. Additionally, viewing **X** through its singular value decomposition (SVD), the PGS vector is a weighted sum of the left singular vectors. Recall that the SVD is **X** = **U** diag(*s*_1_, …, *s*_*r*_) **V**^*T*^, where *r* = min{*n, p*}, **U** = [***u***_1_, …, ***u***_*r*_] is the *n*-by-*r* matrix of left singular vectors, *s*_1_ ≥ *s*_2_ ≥ · · · ≥ *s*_*r*_ ≥ 0 are the singular values, and **V** = [***v***_1_, …, ***v***_*r*_] is the *p*-by-*r* matrix of right singular vectors (principal directions). The *k*th principal component is given by **PC**_*k*_ = *s*_*k*_***u***_*k*_. We collect these statements into Proposition 1 below.

## Proposition 1

(Polygenic Scores are Linear Combinations of PCs and Singular Vectors). *Let* ***β*** *be a length p polygenic effect vector, so that β*_*j*_ *encodes the effect of variant j. Then, the PGS vector* 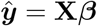 *is a weighted sum of the PCs of* **X**:

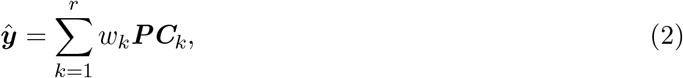

*where the weight w*_*k*_ *associated with* ***PC***_*k*_ *measures (i*.*e*., *is proportional to) the cosine similarity between the kth principal direction* ***v***_*k*_ *and the polygenic effect vector* ***β***. *The PGS vector is also a weighted sum of the left singular vectors of* **X**:

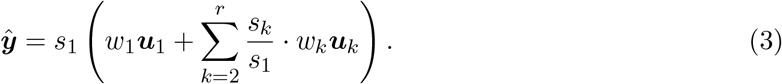

We prove Proposition 1 in Supplementary Material S1. Proposition 1 provides intuition about how a PGS can be evaluated for potential bias by population structure. According to Eq. (2), the empirical distribution of the weights *w*_*k*_ reflects the extent to which a particular PC drives the overall distribution of the PGS vector. Because it is well-recognized that PCs capture population structure (41; 42; 43), Eq. (2) suggests inspecting the empirical distribution of *w*_*k*_ for the training cohort to detect signs of “population structure”-related bias of the polygenic effect vector ***β***. This is closely related to the downstream diagnostic reported in Sohail et al. (34), where the authors first estimated SNP loadings for each PC (equivalent to the principal directions, or the right singular vectors ***v***_*k*_), and then computed Spearman correlations of each loading vector, corresponding to one of the top 40 PCs, with the polygenic effect vector (Supplementary Figure S1A). The Spearman correlation computed for the *k*th PC is analogous to *w*_*k*_ in Eq. (2), and would be in full agreement with our framework had the authors computed cosine similarities between the loading vectors and the polygenic effect vector (see Eq. (S4) in Supplementary Material S1 for exact formula of *w*_*k*_). Nevertheless, the authors’ calculations are tantamount to measuring the relative weight placed on each PC by the PGS, a consequence of the fact that PGS vector can be viewed as a weighted sum of the orthogonal PCs.

Proposition 1 also provides intuition on how a phenotype’s stratification by PCs potentially inflates the performance of a random PGS against the phenotype itself. According to Eq. (3), the overall distribution of the PGS vector is influenced unequally by the collection of singular values (*s*_*k*_ : *k* = 1, …, *r*) of the genotype matrix. In practice, the singular values tend to be dominated by the leading terms (e.g., *s*_1_), a consequence of the empirical observation that the eigenvalues of the sample covariance matrix 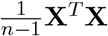 tend to be “spiked” (31; 32); see Supplementary Figure S1B for an example. As a result, many of the terms *s*_*k*_*/s*_1_ appearing within the summands of Eq. (3) are much less than 1. Now, suppose ***β*** is drawn from the standard multivariate normal distribution to construct a random PGS: ***ŷ*** = **X*β***, where ***β*** ∼ 𝒩(**0, I**_*p×p*_). The corresponding weights (*w*_*k*_ : *k* = 1, …, *r*) also follow a standard multivariate normal distribution (see Theorem S1 in Supplement Material). Because each *w*_*k*_ contributes the same effect on average owing to the distributional symmetry, Eq. (3) implies that individual variant effects behaving like random noise can nonetheless produce PGSs that are driven largely by the singular vectors corresponding to the leading singular values. If, additionally, the phenotype were biased by the same PCs that correspond to the leading singular values, then we would expect an inflated performance of such a random projection.

In Supplementary Material S3, we derive an angular central Gaussian null distribution for the vector of magnitudes of the projections of a random projection onto each PC-subspace (Theorem S2). We also report mathematical formulas encoding the impact of a phenotype’s PC and singular value bias on PGS performance, including the widely used phenotype-PGS correlation (Theorem S3) and new performance measures like phenotype-PGS cosine similarity (Theorem S4).

As described in the Results section, our mathematical framework motivates our experiments on UK Biobank (UKB) data. To facilitate reporting the rest of our data processing, we refer to the experiment evaluating performance inflation of rPGSs as **Performance Inflation by rPGS**, and the experiment evaluating the relative performance of original PGSs to random projections as **Performance Relative to pPGS and sPGS**. We refer to our first experiment evaluating MCH PGSs as **Evaluation of MCH PGSs**.

### PGS Catalogue Processing

To obtain counts of variants included in each polygenic score, we accessed raw .txt files following instructions on the REST API page (https://www.pgscatalog.org/rest/#/, date of access: Aug 5, 2023). Because some variants in a score may have an effect size of 0, we did not use the quantity reported in the “Number of Variants” cell under Score Details. Instead, we filtered out variants with zero effect and reported counts of all remaining variants in the score file. A table summarizing non-zero variant counts for all scores is provided on Github, as is the script used for data mining.

### Cohorts and Quantitative Traits

We use a set of *n* = 487, 296 individuals from UKB, most of whom are of self-reported European ancestry. For **Performance Inflation by rPGS**, mirroring the approach of Fiziev and McRae et al. (44) we choose 141 quantitative anthropometric (e.g., height, weight, body fat percentage), blood biomarker (e.g., glucose and LDL concentrations) and urine biomarker (e.g., creatinine) phenotypes. First, to be sufficiently powered, we require that each quantitative phenotype is measured in at least 100, 000 individuals (this excluded all “second visit” phenotypes). We further exclude testosterone, which is a highly sex-specific phenotype. For systolic and diastolic blood pressure, we take the average of the two consecutive measurements at each visit. Each phenotype is then inverse-rank normal transformed (IRNT) and further corrected for a number of other covariates in a phenotype-specific manner, as described in Table S3 (Supplemental Data) of Fiziev and McRae et al. (44). On top of these IRNT phenotypes, the corresponding original version of each phenotype is included in our analyses. This resulted in a total of 141 + 141 = 282 phenotypes. We analyze the Original and IRNT phenotypes separately. The list of quantitative traits is available on Github (see Data Availability).

For **Performance Relative to pPGS and sPGS**, we restrict our cohort to only individuals of self-reported European ancestry (*n*_Eur_ = 357, 659). We chose to focus on individuals of European ancestry, to investigate if PGSs derived from an ancestrally more homogeneous population would still be at risk of population stratification confounding — an issue that has been reported in the literature (9). We further randomly split our restricted cohort, arriving at a training cohort (*n*_train_ = 288, 728) and a held-out test cohort (*n*_test_ = 68, 931), which are used for training PGSs and performing sensitivity analyses respectively. Next, we pick a subset of the 141 phenotypes, based on the number of variants in the trained PGS (see Polygenic Score Construction below for details on our PGS training). Specifically, for inclusion in our analyses, we require the phenotype’s PGS to have at least 5 variants with GWAS *p*-value exceeding the largest perturbation cutoff (10^−6^ for lenient PGS and 10^−10^ for stringent PGS). This ensures that enough distinct perturbed PGSs can be constructed for our sensitivity analyses. For lenient PGSs, this reduces the number of phenotypes to 103; for stringent PGSs, the number is reduced to 68.

Finally, for investigation of mean corpuscular haemoglobin (MCH) PGSs obtained from the PGS Catalogue, we perform all analyses on the held-out test cohort (*n*_test_ = 68, 931).

### Association Mapping

We perform genome-wide association mapping as part of our PGS construction procedure for **Performance Relative to pPGS and sPGS**, as well as for prioritizing variants for performing perturbations in investigating MCH PGSs. Details of quality control (QC) for variants in preparation for association mapping can be found in Churchhouse et al. (45). Briefly, our genome-wide association study (GWAS) cohort comprises 357, 661 unrelated individuals of selfreported white British ancestry. We excluded non-European participants to reduce confounding due to population structure. A total of 13, 791, 468 genotyped common variants (AF *>* 0.01) are used, and are either directly measured or imputed across the individuals. The individuals are restricted to have self-declared white British genetic ancestry, with exclusion of closely related individuals (or at least one of a related pair of individuals), individuals with sex chromosome aneuploidies, and individuals who had withdrawn consent from the UKB study.

We perform GWAS on quantitative phenotypes using imputed genotypes from the UK Biobank. For each trait we include the 13.8 million variants that passed QC. Each variant is tested for association with the trait under an additive model, using the dosage of the minor allele. Analysis of each trait used phenotype values which had been adjusted for age, sex, and usage of relevant medications, and were inverse rank normalized prior to analysis. The marginal significance of each variant was tested with a linear model which included age, sex, and the first ten genetic principal components as covariates. Details on Principal Components Analysis are reported in Supplementary Material Section S3 of (46). Covariates are standardized to mean = 0 and standard deviation = 1 before regression.

### Polygenic Score Construction and Processing

For **Performance Inflation by rPGS**, each random PGS (rPGS) is independently constructed by selecting 10% of all available autosomal variants uniformly at random (940, 851 variants selected each time); effect sizes are independently drawn from the standard Gaussian distribution. For **Performance Relative to pPGS and sPGS**, PGSs are constructed using Clumping and Thresholding (C&T) (47). Variants within a 1Mb window of a GWAS peak are clumped. Further, peaks out to 10Mb in high LD (i.e., *r*^2^ *>* 0.1) are excluded in order to avoid re-including non-independent peaks and to ensure windows were non-overlapping. During stepwise selection of variants within a 1Mb window, a restriction of three variants per window is applied. The final PGS is obtained through an effect size re-weighting step: multiple regressions were performed on the selected variants within each window, before aggregating the resulting reweighted effects across all windows. We explore various *p*-value thresholds (from 10^−100^ to 10^−4^, see Figure 1A), and choose 10^−5^ and 10^−8^ as two thresholds that result in two PGSs constructed per phenotype. We refer to the former PGS as the lenient PGS, which includes variants that do not pass genome-wide significance but on average is more performant, as measured by squared correlation coefficient. The latter PGS is referred to as the stringent PGS. PGS files are available on Github (see Data Availability).

For our investigation of MCH PGSs, owing to differences in training cohorts and the lack of information regarding each variant’s significance in the publicly-available PGS Catalogue files, we match variants appearing in each PGS with our GWAS summary statistics. The average fraction of unmatched variants across PGSs is 9%, with a majority of PGSs achieving at least a 97% matching fraction (Supplementary Table S5). For PGSs that contain multiple variants on the same position, we kept the variant that had the smallest GWAS *p*-value, or picked one at random if the *p*-values were unavailable. Such variants, on average, accounted for less than 0.035% of variants included in PGSs.

We perform computations in R. To address the challenge of high-dimensional matrix-vector operations associated with PGS computations, we compute all PGSs using bigsnpr package (48).

### PGS Relative Performance Evaluation

When evaluating the relative performance of an original PGS, we construct 200 random projections (100 pPGSs and 100 sPGSs). For each random projection ***ŷ*** (either a pPGS or a sPGS), we compute its performance against the original phenotype ***y***, based on a chosen performance metric, as follows. For a fixed choice of perturbation threshold parameter *p*_non-sig_ (we consider four values, 10^−6^, 10^−7^, 10^−8^ and 10^−10^, altogether across lenient and stringent PGSs), the set of variants included in the PGS is split into two subsets, one corresponding to variants whose GWAS *p*-value exceeds *p*_non-sig_ (i.e., non-significant) and whose effects will thus be drawn at random, and the other corresponding to variants whose *p*-value is smaller than *p*_non-sig_ (i.e., their *β*_*j*_’s are fixed throughout). By summing the corresponding effects across variants from each subset, one obtains two scores, ***ŷ***_*R*_ and ***ŷ***_*S*_, that sum to the PGS vector: ***ŷ*** = ***ŷ***_*S*_ + ***ŷ***_*R*_. (Note the subscripts *S* and *R* denote “Significant” and “Random,” respectively.) Concretely, ***ŷ***_*R*_ = Σ_variant *j* in subset 1_ *β*_*j*_ ***x***^(*j*)^ and ***ŷ***_*S*_ = Σ_variant *j* in subset 2_ *β*_*j*_ ***x***^(*j*)^, with ***x***^(*j*)^ denoting the *j*th column of genotype matrix **X**. Viewing the effects of variants contributing to ***ŷ***_*R*_ as random, we additionally compute a “reflected” PGS, ***ŷ***_−_ = ***ŷ***_*S*_ − ***ŷ***_*R*_. Now let *f* = *f* (***ŷ, y***) be a given metric, for example, *f* could be the rank correlation between ***ŷ*** and ***y***. We compute two quantities, *f* ^+^ := *f* (***ŷ, y***) and *f* ^−^ := *f* (***ŷ***_−_, ***y***). Finally, the performance is reported as max{*f* ^+^, *f* ^−^}, to account for the fact that random effects have equal chances of being positively or negatively correlated with the phenotype (see Supplementary Material S8 for details). In **Evaluation of MCH PGSs**, we used *p*_non-sig_ = 10^−6^ to analyze and report our results. However, in general, on top of using multiple *p*_non-sig_’s as reported in Results, we also evaluated PGSs the usual way; that is, defining performance by *f* (***ŷ, y***) instead of max{*f* ^+^, *f* ^−^}, and performing all analyses with respect to such an approach. We found that the results did not change qualitatively (details reported in Supplementary Material S8).

### Evaluating Statistical Significance of Relationships

We perform various statistical tests to identify significant relationships (e.g., two-sample comparisons for detecting dominance of one distribution over another, correlations for detecting trends relating two quantities). Throughout this work, We use non-parametric and distribution-free tests (e.g., Wilcoxon rank sum test and rank correlation test), to achieve Type I Error control. Because of the multiplicity of metrics and measures considered in our work, we further perform Bonferroni correction to control the familywise error rate at 0.05.

