## Supplementary Material for "Highly parameterized polygenic scores tend to overfit to population stratification via random effects"

##### S1 Proofs of Main Results

In this Section we prove the main mathematical result reported in the Main Text. We also quickly state and prove a claim reported in the Main Text, to do with the weights of the linear combination representation of a PGS following a standard multivariate Gaussian distribution whenever the effects follow the standard multivariate Gaussian distribution.

*Proof of Proposition 1.* First, we shall show that a vector of polygenic scores — defined as the matrix-by-vector product between the cohort's normalized genotype matrix and the polygenic weight vector — can be expressed as a linear combination of the principal components of that

cohort; that is, it obeys Eq. (2) in Proposition 1. Formally, let

$$\mathbf{X} = \begin{pmatrix} — & \boldsymbol{x}_1 & — \\ — & \vdots & — \\ — & \boldsymbol{x}_n & — \end{pmatrix} \in \mathbb{R}^{n\times p}$$

denote the normalized allelic dosage matrix of $n$ individuals across $p$ variants. Normalized means that the $j$th column of the allelic dosage matrix has $2\widehat{f}_j$ subtracted from it before being scaled down by $\sqrt{2\widehat{f}_j(1-\widehat{f}_j)}$ where $\widehat{f}_j$ is the cohort's allele frequency. Let $\boldsymbol{\beta} \in \mathbb{R}^p$ denote the effect vector.

Let $r = \min\{n, p\}$. Recall the singular value decomposition (SVD) of $\mathbf{X}$ allows the genotype matrix to be written as a sum of rank one matrices:

$$\begin{aligned} \mathbf{X} &= \underset{n\times r}{\mathbf{U}} \underset{r\times r}{\mathrm{diag}(s_k)} \underset{r\times p}{\mathbf{V}^T} \\ &= \begin{pmatrix} | & \cdots & | \\ \boldsymbol{u}_1 & \cdots & \boldsymbol{u}_r \\ | & \cdots & | \end{pmatrix} \begin{pmatrix} s_1 & \cdots & 0 \\ \vdots & \ddots & \vdots \\ 0 & \cdots & s_r \end{pmatrix} \begin{pmatrix} — & \boldsymbol{v}_1^T & — \\ — & \vdots & — \\ — & \boldsymbol{v}_r^T & — \end{pmatrix} \\ &= \sum_{k=1}^{r} s_k \boldsymbol{u}_k \boldsymbol{v}_k^T. \end{aligned} \tag{S1}$$

Moreover, we have the following relationship between the left singular vectors and the right singular vectors: for all $k \in [r]$,

$$\boldsymbol{u}_k = \frac{1}{s_k}\mathbf{X}\boldsymbol{v}_k. \tag{S2}$$

The final ingredient we need to verify Eq. (2) is the eigendecomposition formula that is central to principal components analysis (PCA). Given $\mathbf{X}$, which is column-centered, we may construct its covariance matrix $\mathbf{C} = \frac{1}{n-1}\mathbf{X}^T\mathbf{X}$. Then, by the SVD we see that

$$\mathbf{C} = \frac{1}{n-1}(\mathbf{U}\ \mathrm{diag}(s_k)\ \mathbf{V}^T)^T(\mathbf{U}\ \mathrm{diag}(s_k)\ \mathbf{V}^T) = \mathbf{V}\ \mathrm{diag}\left(\frac{s_k^2}{n-1}\right)\ \mathbf{V}^T.$$

The column vectors $(\boldsymbol{v}_k : k = 1, \ldots, r)$ that make up $\mathbf{V}$ are the *principal directions*, and *principal components* (PCs) are obtained by right-multiplying the matrix $\mathbf{X}$ by the corresponding principal directions: that is, for all $k \in [r]$,

$$\boldsymbol{PC}_k = \mathbf{X}\boldsymbol{v}_k. \tag{S3}$$

We can now work through the steps to obtain Eq. (2). Starting with a vector of effect sizes $\boldsymbol{\beta}$, possibly obtained from fitting a polygenic score training algorithm on $\mathbf{X}$ itself,

$$\begin{aligned} \mathbf{X}\boldsymbol{\beta} &= \left(\sum_{k=1}^{r} s_k \boldsymbol{u}_k \boldsymbol{v}_k^T\right)\boldsymbol{\beta} \\ &= \sum_{k=1}^{r} s_k \boldsymbol{u}_k \langle \boldsymbol{v}_k, \boldsymbol{\beta}\rangle \qquad (*) \\ &= \sum_{k=1}^{r} \mathbf{X}\boldsymbol{v}_k \cdot \langle \boldsymbol{v}_k, \boldsymbol{\beta}\rangle \quad \text{(by Eq. (S2))} \\ &= \sum_{k=1}^{r} \boldsymbol{PC}_k \cdot w_k, \qquad \text{(by Eq. (S3))} \end{aligned}$$

where

$$w_k = \langle \boldsymbol{v}_k, \boldsymbol{\beta} \rangle. \tag{S4}$$

Thus, Eq. (2) is verified.

Moreover, in the steps above, simply by replacing $\langle \boldsymbol{v}_k, \boldsymbol{\beta} \rangle$ with $w_k$ in $(*)$, we can easily arrive at Eq. (3) of Proposition 1. This verifies Eq. (3).

To conclude our proof, we note that the quantity $w_k$ has a convenient interpretation: it measures the similarity of the effect size vector to the $j$th principal direction. Indeed, from the definition of the inner product, we see that

$$w_k = ||\boldsymbol{v}_k||_2 ||\boldsymbol{\beta}||_2 \cos(\theta_k) = ||\boldsymbol{\beta}||_2 \cos(\theta_k),$$

where $\theta_k$ measures the angle between $\boldsymbol{v}_k$ and $\boldsymbol{\beta}$. □

In the Materials and Methods section of our Main Text, we claim that the distribution of the vector of weights $\boldsymbol{w} = (w_k : k = 1, \ldots, r)$ is a standard multivariate normal whenever the polygenic weight vector $\boldsymbol{\beta}$ itself is drawn from a standard multivariate normal distribution. We state this claim precisely below and provide a quick proof.

**Theorem S1.** *Let $\boldsymbol{w} = (w_k : k = 1, \ldots, r)$ be the cosine similarity weights defined in Eq.* (S4)*. Suppose that the polygenic weight vector $\boldsymbol{\beta}$ follows a standard multivariate Gaussian distribution: $\beta_j \overset{iid}{\sim} \mathcal{N}(0,1)$. Then $\boldsymbol{w}$ follows a standard multivariate Gaussian distribution: $w_k \overset{iid}{\sim} \mathcal{N}(0,1)$.*

*Proof of Theorem S1.* We may write

$$\boldsymbol{w} = \begin{pmatrix} — & \boldsymbol{v}_1^T & — \\ — & \vdots & — \\ — & \boldsymbol{v}_r^T & — \end{pmatrix} \boldsymbol{\beta} = \mathbf{V}^T \boldsymbol{\beta}. \tag{S5}$$

Because $\boldsymbol{\beta} \sim \mathcal{N}(\mathbf{0}, \mathbf{I}_{p\times p})$, by affine properties of the multivariate Gaussian,

$$\begin{aligned} \boldsymbol{w} &\sim \mathcal{N}\left(\mathbf{V}^T \mathbb{E}[\boldsymbol{\beta}], \mathbf{V}^T \mathrm{Cov}(\boldsymbol{\beta}) \mathbf{V}\right) \\ &= \mathcal{N}(\mathbf{0}, \mathbf{I}_{r\times r}) \end{aligned}$$

where the first line follows from Eq. (S5) and the second line follows from orthogonality of $\mathbf{V}$ and the fact that $\boldsymbol{\beta}$ has the standard multivariate normal distribution. □

As a pedantic remark, if $\mathbf{X}$ is not full rank — i.e., some of the singular values in the SVD are 0 — then the result in Theorem S1 still holds for all indexes $k$ as long as the corresponding singular value $s_k$ is non-zero. This scenario occurs, for example, if the columns of $\mathbf{X}$ are multicollinear and there are fewer variants $p$ than individuals $n$.

#### S2 Violation of Simplifying Assumptions

Our mathematical results and their accompanying proofs rely on two simplifying assumptions, which we state in the Main Text but also include below for convenience.

1. The set of $p$ variants making up $\mathbf{X}$ are those used in constructing the PCs, and these variants moreover include all variants selected for inclusion during PGS training.

2. The PGS is computed on the normalized version of $\mathbf{X}$, on which PCA was previously performed. That is, $\mathbf{X}$ is obtained from the original genotype matrix of allelic dosages, by subtracting the $j$th column by $2\widehat{f}_j$ before scaling it down by $\sqrt{2\widehat{f}_j(1-\widehat{f}_j)}$ where $\widehat{f}_j$ is the cohort's (estimated) allele frequency. Remark that the factor of 2 in the last quantity is for variance standardization. The implicit assumption is that the cohort is in panmixia and at equilibrium, with diploid allelic dosages assigned to each individual according to the Binomial model, $\text{Binom}(2, f_j)$, where $f_j$ is unknown and hence estimated by $\hat{f}_j$. This is consistent with how the software PLINK (Purcell et al., 2007) computes genetic PCs from the genetic relatedness matrix as defined in Yang et al. (2011).

Both assumptions are violated in practice. Regarding Assumption 1, PCA is typically performed on an LD-pruned version of the genotype matrix, amongst other technical processing steps. For instance, in our present work, the set of variants was pruned based on LD threshold $r^2 < 0.1$, using windows of 1000 base pairs (bp) and a step-size of 80bp. (All other technical processing steps are detailed in pp. 17-18 in the Supplementary Material file of Bycroft et al., 2017.) Variant inclusion criteria for a PGS may use different LD thresholds and window sizes, or may even ignore linkage concerns (e.g., LDPred2 with the infinitesimal prior setting, Privé et al., 2020).

Regarding Assumption 2, PGSs are computed on the allelic dosage matrix, with the effect $\beta_j$ encoding the marginal increase in the polygenic score of an individual per unit increase in allelic dosage. While this technical detail obscures the clarity of presentation of our mathematical results, it is possible to work out the exact relationship between the PGS vector and the PCs under this more realistic setting. We do so below.

Let $\widehat{f}_j$ denote the estimated allele frequency of variant $j$ for a cohort. Let $\mathbf{X}$ denote the allelic dosage matrix of the cohort, and suppose that $\mathbf{X}'$ is the normalized version of it. Concretely, for column $j$ of $\mathbf{X}$, subtract $2\widehat{f}_j$ from each entry before scaling it down by $\sqrt{2\widehat{f}_j(1-\widehat{f}_j)}$. The resulting column vector is the $j$th column of $\mathbf{X}'$. Now let $\boldsymbol{\beta}$ denote the polygenic effect vector, where $\beta_j$ is the marginal increase in polygenic score per unit increase in allelic dosage. Define an allele-frequency-scaled version of $\boldsymbol{\beta}$, which we write as $\boldsymbol{\beta}'$, by multiplying the effect of the $j$th variant by the Hardy-Weinberg standard deviation factor: $\beta'_j = \beta_j \cdot \sqrt{2\widehat{f}_j(1-\widehat{f}_j)}$. It is straightforward to show that

$$\mathbf{X}\boldsymbol{\beta} = \mathbf{X}'\boldsymbol{\beta}' + \left(\sum_{j=1}^{p} 2\widehat{f}_j\beta_j\right)\mathbf{1}, \tag{S6}$$

where $\mathbf{1}$ denotes the length $p$ vector of ones. From Eq. (S6) we see that the PGS vector (LHS) and its allele-frequency-scaled analogue differ by a constant term, which is determined by a weighted sum of genetic effects with weights corresponding to the cohort allele frequencies (RHS). To obtain the desired relationship between the PGS vector and the genetic PCs of $\mathbf{X}'$, as is done in practice, we apply the SVD argument in the proof of Proposition 1 to $\mathbf{X}'$. This results in the PGS vector $\hat{\boldsymbol{y}}$ satisfying

$$\begin{aligned} \hat{\boldsymbol{y}} &= \mathbf{X}\boldsymbol{\beta} \\ &= \sum_{k=1}^{r} \boldsymbol{PC}_k \cdot w'_k + \left(\sum_{j=1}^{p} 2\widehat{f}_j\beta_j\right)\mathbf{1}, \end{aligned} \tag{S7}$$

where $w'_k = ||\boldsymbol{\beta}'||_2 \cos(\theta'_k)$, where $\theta'_k$ measures the angle between the $k$th right singular vector of $\mathbf{X}'$ and $\boldsymbol{\beta}'$. Therefore, as Eq. (S7) shows, in practice the PGS vector is the sum of a linear combination of PCs and a constant term involving estimated allele frequencies. Owing to little conceptual and empirical clarity gained (in our present work) from working with Eq. (S7), in our presentation of results we have chosen to stick with Assumption 2.

#### S3 Consequences of Stratification

In this Section we apply our framework and Proposition 1 to concretely describe the impact of population structure and PC stratification on the behaviour of a PGS. First, we derive a null distribution of the projection of a random PGS onto the subspace spanned by the $k$th PC. This result reveals a mathematical relationship between the singular values of the genotype matrix and the "degree of alignment" of a random PGS with each PC computed on the same genotype matrix.

Recall that the *Angular Central Gaussian (ACG) distribution* is a distribution parameterized by a positive semidefinite matrix $\mathbf{A}$ and supported on the $(r-1)$-dimensional hypersphere $\mathbb{S}^{r-1} = \{\boldsymbol{\ell} \in \mathbb{R}^r : ||\boldsymbol{\ell}||_2 = 1\}$ (Tyler, 1987; see also Section 9.4.4 on pp. 182-183 of Mardia and Jupp, 2000). The density function is given by

$$f(\boldsymbol{\ell};\mathbf{A}) = \alpha_r^{-1}\det(\mathbf{A})^{-\frac{1}{2}}(\boldsymbol{\ell}^T\mathbf{A}^{-1}\boldsymbol{\ell})^{-\frac{r}{2}}, \quad \text{(S8)}$$

where $\alpha_r = 2\pi^{\frac{r}{2}}/\Gamma(\frac{r}{2})$ is the surface area of $\mathbb{S}^{r-1}$ and $\Gamma(x)$ is the Euler gamma function (Abramowitz et al., 1988). We write $\boldsymbol{\ell} \sim ACG(\mathbf{A})$ whenever the density function of the random vector $\boldsymbol{\ell}$ is given by Eq. (S8). Additionally, the following properties hold of the ACG distribution.

**Property 1.** The ACG distribution arises from projecting a multivariate Gaussian onto the hypersphere. That is, if $\boldsymbol{z} \sim \mathcal{N}(\mathbf{0},\mathbf{A})$, then $\boldsymbol{z}/||\boldsymbol{z}||_2 \sim ACG(\mathbf{A})$.

**Property 2.** The ACG distribution has a degree of indeterminacy. Specifically, for any pair of positive semidefinite matrices $\{\mathbf{A},\mathbf{B}\}$ satisfying $\mathbf{A} = c\mathbf{B}$ for some positive scalar $c$, $ACG(\mathbf{A}) \overset{d}{=} ACG(\mathbf{B})$ (i.e., the distributions are equal).

**Property 3.** The ACG distribution has simple moments in the standard multivariate Gaussian case. That is, if $\mathbf{A} = \mathbf{I}_{r\times r}$, then the following quantities describe the analytical moments of $ACG(\mathbf{A})$:

$$\begin{aligned} \mathbb{E}[\boldsymbol{\ell}] &= \mathbf{0} \\ \mathrm{Cov}(\boldsymbol{\ell}) &= \mathrm{diag}(1/r) \end{aligned}$$

Formulas for higher moments are described in Section 9.6.1 of Mardia and Jupp (2000).

These properties are useful for deriving and interpreting the following result.

**Theorem S2** (Null Distribution of PC-Projection of Random PGS)**.** *Let* $\ell_k = \hat{\boldsymbol{y}}^T\boldsymbol{PC}_k/(||\hat{\boldsymbol{y}}||_2||\boldsymbol{PC}_k||_2)$ *denote the normalized projection of the PGS vector* $\hat{\boldsymbol{y}}$ *onto the subspace spanned by the kth PC, where* $\hat{\boldsymbol{y}} = \mathbf{X}\boldsymbol{\beta}$ *is obtained by a standard multivariate Gaussian polygenic weight vector* $\boldsymbol{\beta}$ *as in Theorem S1. Then, the random vector* $\boldsymbol{\ell} = (\ell_1,\ldots,\ell_r)$ *satisfies*

$$\boldsymbol{\ell} \sim ACG(diag(s_1^2,\ldots,s_r^2)), \quad \text{(S9)}$$

*where* $s_k$ *is the kth singular value as defined in Proposition 1 in the Main Text.*

*Proof.* Since $\boldsymbol{PC}_k = s_k\boldsymbol{u}_k$ (see eqs. (S2) and (S3)), applying the invariance to scaling of the normalized projection, we obtain $\ell_k = \hat{\boldsymbol{y}}^T\boldsymbol{u}_k/(||\hat{\boldsymbol{y}}||_2||\boldsymbol{u}_k||_2) = \hat{\boldsymbol{y}}^T\boldsymbol{u}_k/||\hat{\boldsymbol{y}}||_2$. By using Eq. (3) of Proposition 1 twice, we obtain the two equations,

$$\begin{aligned}\hat{\boldsymbol{y}}^T\boldsymbol{u}_k &= s_k w_k,\\ ||\hat{\boldsymbol{y}}||_2 &= s_1^2w_1^2 + \ldots + s_r^2 w_r^2.\end{aligned}$$

Defining $z_k = s_k w_k$, it is clear from $w_k \overset{\text{iid}}{\sim} \mathcal{N}(0,1)$ — which we proved in Theorem S1 earlier — that the random vector $\boldsymbol{z} = (z_1,\ldots,z_r) \sim \mathcal{N}(\boldsymbol{0}, \text{diag}(s_1^2,\ldots,s_r^2))$, and that $\boldsymbol{\ell} = \boldsymbol{z}/||\boldsymbol{z}||_2$. Applying Property 1 with $\mathbf{A} = \text{diag}(s_1^2,\ldots,s_r^2))$ finishes the proof. □

We remark that $\ell_k$ as defined in Theorem S2 is the cosine similarity of the PGS vector with the $k$th PC. Now let us illustrate how Eq. (S9) captures the impact of population structure on the behaviour of a random PGS. In the case where the singular values are all equal (we may set them all to 1 by Property 2), Property 3 shows that the projections of a random PGS onto the PC subspaces are uncorrelated, and moreover the variances — which measure the magnitudes of the projections — are all equal to $1/r$; see Figure S2A. On the other hand, when the singular values are unequal, as is typical of genotype matrices, the behaviour of the projections onto PC subspaces will no longer agree with the equations in Property 3. Instead, their behaviour follows from the following property.

**Property 4.** If $\mathbf{A} = \text{diag}(s_1^2,\ldots,s_r^2)$, then the mean of $ACG(\mathbf{A})$ is $\mathbb{E}[\boldsymbol{\ell}] = \boldsymbol{0}$, while the covariance is $\text{Cov}(\boldsymbol{\ell}) = \mathbb{E}\left[\frac{\mathbf{A}^{1/2}\boldsymbol{z}\boldsymbol{z}^T\mathbf{A}^{1/2}}{\boldsymbol{z}^T\mathbf{A}\boldsymbol{z}}\right]$, where $\boldsymbol{z} \sim ACG(\mathbf{I}_{r\times r})$.

In other words, although the projections are still 0 on average, the magnitudes of the projections depend on the singular values $s_k$. When $r = 2$, one can rely on a polar reparameterization argument[1] to prove that

$$\text{Cov}(\boldsymbol{\ell}) = \begin{pmatrix}\frac{s_1}{s_1+s_2} & 0\\ 0 & \frac{s_2}{s_1+s_2}\end{pmatrix}.$$

This shows that the "relative degree of alignment" of the PGS with the top PC (i.e., $\text{Var}(\ell_1)/\text{Var}(\ell_2)$) is determined precisely by the ratio of the singular values $s_1/s_2$. For $r \geqslant 3$, it is more challenging to obtain the analytical expression of the analogous covariance matrix. However, by performing a simple simulation[2] using the singular values reported in Supplementary Table S2 of Galinsky et al. (2016) (we assume here that $r = 10$ because only 10 eigenvalues were reported), we observe empirically the diagonal of $\text{Cov}(\boldsymbol{\ell})$ follows a more skewed vector than the analytical generalization of the $r = 2$ case, which is $\left(\frac{s_1}{s_1+\ldots+s_r},\ldots,\frac{s_r}{s_1+\ldots+s_r}\right)$; see Figure S2B. This suggests that the relative degree of alignment of the PGS with the top PC may be more skewed than the ratio of the singular values when more than two variants are included, which is the case for the majority (if not all) of Biobank-scale polygenic analyses.

Next, we demonstrate mathematically how stratification by PCs may introduce potential bias to evaluation metrics used to diagnose PGS efficacy. We show how the Pearson correlation between a PGS and the phenotype is related to (1) the weights $w_k$ capturing alignment of the polygenic effect vector $\boldsymbol{\beta}$ with the $k$th PC, (2) the singular values of the cohort genotype matrix, $\boldsymbol{s} = (s_1,\ldots,s_r)$,

[1]See `https://stats.stackexchange.com/questions/616404/non-uniform-spherical-distributions/`.

[2]Briefly, we independently draw $R = 10^4$ vectors, $\boldsymbol{\ell}_1,\ldots,\boldsymbol{\ell}_R$, from $ACG(\text{diag}(s_1^2,\ldots,s_r^2))$, before computing the empirical mean squared value of each component to estimate the second moment. Simulations are performed in **R** using `rotasym` package (García-Portugués et al., 2020), and code is available on Github at: `https://github.com/songlab-cal/StratPGS`.

and (3) the empirical correlations between the phenotype and each PC. We also show how the cosine similarity between a PGS and the phenotype is related to (1) the weights $w_k$ capturing alignment of the polygenic effect vector $\boldsymbol{\beta}$ with the $k$th PC, (2) the singular values of the cohort genotype matrix, $\boldsymbol{s} = (s_1, \ldots, s_r)$, and (3) the empirical cosine similarities between the phenotype and each PC. We first state and prove these results; a brief discussion is included afterward.

**Theorem S3** (PGS-Phenotype Correlation)**.** *Let the phenotype vector be $\boldsymbol{y}$, and suppose that its empirical correlation with the $k$th principal component of the training cohort is $r_k = Corr(\boldsymbol{y}, \boldsymbol{PC}_k)$, where $\boldsymbol{PC}_k$ is computed from the $n \times p$ genotype matrix $\mathbf{X}$. Let $(r_k : k = 1, \ldots, r)$ be the collection of empirical correlations. Suppose, as in Eq.* (2) *of Proposition 1, that for a given PGS $\hat{\boldsymbol{y}}$ and polygenic weight vector $\boldsymbol{\beta}$, the associated weights from projecting $\boldsymbol{\beta}$ onto PC subspaces are $\boldsymbol{w} = (w_1, \ldots, w_r)$. Suppose further, as in Eq.* (3) *of Proposition 1, that the singular values and left singular vectors associated with $\mathbf{X}$ are $\boldsymbol{s} = (s_1, \ldots, s_r)$ and $(\boldsymbol{u}_1, \ldots, \boldsymbol{u}_r)$, respectively. For any cohort size $n$,*

$$Corr(\boldsymbol{y}, \hat{\boldsymbol{y}}) = \sum_{k=1}^{r} \frac{w_k r_k s_k \sqrt{1 - n\mu_{\boldsymbol{u}_k}^2}}{\sqrt{w_1^2 s_1^2 + \ldots + w_r^2 s_r^2 - n\mu_{\hat{\boldsymbol{y}}}^2}} \tag{S10}$$

*where we use $\mu_{\boldsymbol{v}}$ to denote the mean of a vector $\boldsymbol{v}$.*

*Proof.* First, observe that $r_k = \mathrm{Corr}(\boldsymbol{y}, \boldsymbol{PC}_k)$ implies that

$$(\boldsymbol{y} - \mu_{\boldsymbol{y}}\mathbf{1})^T(\boldsymbol{PC}_k - \mu_{\boldsymbol{PC}_k}\mathbf{1}) = r_k||\boldsymbol{y} - \mu_{\boldsymbol{y}}\mathbf{1}||_2||\boldsymbol{PC}_k - \mu_{\boldsymbol{PC}_k}\mathbf{1}||_2, \tag{S11}$$

where $\mu_{\boldsymbol{v}}$ denotes the mean of all elements of a vector $\boldsymbol{v}$. Next, observe that $\hat{\boldsymbol{y}} = \mathbf{X}\boldsymbol{\beta} = \sum_{k=1}^{r} w_k \boldsymbol{PC}_k$ implies that

$$\hat{\boldsymbol{y}} - \mu_{\hat{\boldsymbol{y}}}\mathbf{1} = \sum_{k=1}^{r} w_k \left(\boldsymbol{PC}_k - \mu_{\boldsymbol{PC}_k}\mathbf{1}\right). \tag{S12}$$

Putting together Eqs. (S11) and (S12), we obtain

$$\begin{aligned}(\boldsymbol{y} - \mu_{\boldsymbol{y}}\mathbf{1})^T \left(\hat{\boldsymbol{y}} - \mu_{\hat{\boldsymbol{y}}}\mathbf{1}\right) &= \sum_{k=1}^{r} w_k \left(\boldsymbol{y} - \mu_{\boldsymbol{y}}\mathbf{1}\right)^T \left(\boldsymbol{PC}_k - \mu_{\boldsymbol{PC}_k}\mathbf{1}\right) \\ &= \sum_{k=1}^{r} w_k r_k ||\boldsymbol{y} - \mu_{\boldsymbol{y}}\mathbf{1}||_2 ||\boldsymbol{PC}_k - \mu_{\boldsymbol{PC}_k}\mathbf{1}||_2. \end{aligned} \tag{†}$$

The last piece of information we need is Eq. (S2) from the proof of Proposition 1, which shows that

$$\hat{\boldsymbol{y}} = \sum_{k=1}^{r} w_k s_k \boldsymbol{u}_k, \tag{S13}$$

where $s_k$ and $\boldsymbol{u}_k$ are the $k$th singular value and $k$th left singular vector as defined in Proposition 1.

Putting together the pieces now, we plug Eq. (†) into the definition of correlation, and obtain

$$\begin{aligned}\mathrm{Corr}(\boldsymbol{y}, \hat{\boldsymbol{y}}) &= \frac{\frac{1}{n-1}\left(\boldsymbol{y}-\mu_{\boldsymbol{y}}\mathbf{1}\right)^T\left(\hat{\boldsymbol{y}}-\mu_{\hat{\boldsymbol{y}}}\mathbf{1}\right)}{\frac{1}{n-1}||\boldsymbol{y}-\mu_{\boldsymbol{y}}\mathbf{1}||_2||\hat{\boldsymbol{y}}-\mu_{\hat{\boldsymbol{y}}}\mathbf{1}||_2} \\ &= \sum_{k=1}^{r}\frac{w_k r_k||\boldsymbol{PC}_k-\mu_{\boldsymbol{PC}_k}\mathbf{1}||_2}{||\hat{\boldsymbol{y}}-\mu_{\hat{\boldsymbol{y}}}\mathbf{1}||_2} \\ &= \sum_{k=1}^{r}\frac{w_k r_k s_k||\boldsymbol{u}_k-\mu_{\boldsymbol{u}_k}\mathbf{1}||_2}{\sqrt{w_1^2 s_1^2+\ldots+w_r^2 s_r^2-n\mu_{\hat{\boldsymbol{y}}}^2}} \\ &= \sum_{k=1}^{r}\frac{w_k r_k s_k\sqrt{1-n\mu_{\boldsymbol{u}_k}^2}}{\sqrt{w_1^2 s_1^2+\ldots+w_r^2 s_r^2-n\mu_{\hat{\boldsymbol{y}}}^2}}.\end{aligned}$$

□

**Theorem S4** (PGS-Phenotype Cosine Similarity)**.** *As in Theorem S3 let the phenotype vector be $\boldsymbol{y}$, and suppose that its cosine similarity with the kth principal component of the training cohort is given by $a_k = CosSim(\boldsymbol{y}, \boldsymbol{PC}_k)$. Let $(a_k : k = 1, \ldots, r)$ be the collection of empirical cosine similarities. Suppose, as in Eq. (2) of Proposition 1, that for a given PGS $\hat{\boldsymbol{y}}$ and polygenic weight vector $\boldsymbol{\beta}$, the associated weights from projecting $\boldsymbol{\beta}$ onto PC subspaces are $\boldsymbol{w} = (w_1, \ldots, w_r)$. Suppose further, as in Eq. (3) of Proposition 1, that the singular values associated with $\mathbf{X}$ are $\boldsymbol{s} = (s_1, \ldots, s_r)$. Then,*

$$CosSim(\boldsymbol{y}, \hat{\boldsymbol{y}}) = \sum_{k=1}^{r}\frac{w_k a_k s_k}{\sqrt{w_1^2 s_1^2+\ldots+w_r^2 s_r^2}}. \quad \text{(S14)}$$

*Proof.* The proof strategy is identical to that of Theorem S3, so we sketch the steps of the proof and omit details. Similar to the proof of Theorem S3, observe that $a_k = \mathrm{CosSim}(\boldsymbol{y}, \boldsymbol{PC}_k)$ implies that

$$\boldsymbol{y}^T\boldsymbol{PC}_k = a_k||\boldsymbol{y}||_2||\boldsymbol{PC}_k||_2. \quad \text{(S15)}$$

Now use the fact $\hat{\boldsymbol{y}} = \sum_{k=1}^{r} w_k\boldsymbol{PC}_k$ and Eq. (S15) to simplify the quantity $\boldsymbol{y}^T\hat{\boldsymbol{y}}$. To obtain Eq. (S14) and finish the proof, one just needs to recall that $\mathrm{CosSim}(\boldsymbol{y}, \hat{\boldsymbol{y}}) = \boldsymbol{y}^T\hat{\boldsymbol{y}}/(||\boldsymbol{y}||_2||\hat{\boldsymbol{y}}||_2)$, and divide the quantity obtained in the last step by $||\boldsymbol{y}||_2||\hat{\boldsymbol{y}}||_2$. □

We now discuss the implications of Theorems S3 and S4. The two theorems reveal that the correlation and cosine similarity between any PGS and the phenotype vector depend explicitly on how three empirical sets of quantities — (1) alignment of $\boldsymbol{\beta}$ with each PC (the $w_k$'s), (2) singular values of the genotype matrix (the $s_k$'s), and (3) correlation (resp., cosine similarity) between the phenotype and each PC (the $r_k$'s (resp., $a_k$'s)) — interact in Eq. (S10). In particular, if a PGS were generated from independent random effects — specifically setting $\beta_k$'s to standard normal Gaussians — which would imply that $w_k$'s are independent and identically distributed standard Gaussians (Theorem S1), then the null distribution would still depend on how the leading singular values match with the correlations between the phenotype and the top PCs. While the exact dependencies are unlikely to be worked out analytically, a simulation study would clarify the impact of the three empirical quantities on PGS-phenotype relationships. We defer this study, as well as related investigations into other popular PGS performance metrics, to future work.

#### S4 Metrics and Measures

This Section summarizes all measures and metrics used for quantifying the concepts introduced in our Main Text, including PGS Performance and PC Stratification (see Table 2 of the Main Text).

**Performance Metrics.** To evaluate PGS performance, we computed 11 metrics. We list these metrics in Supplementary Table S1. For **Performance Inflation by rPGS**, we summarize the performance of 200 rPGSs by computing the following summaries of the metrics annotated in Supplementary Table S1.

- Incremental $R^2$: mean
- Percentile-Prevalence Correlation: mean absolute value

**PC Stratification Measures.** To evaluate PC stratification of either a PGS or a phenotype, we computed 12 measures. We list these measures in Supplementary Table S2.

**Perturbed-Fixed Architecture Measures.** In our **Performance Relative to pPGS and sPGS** experiment, we computed multiple measures to quantify distributional properties of the fixed variants and perturbed variants (variants with sign-flipped or permuted effects) of the polygenic effect vector, across multiple perturbation cutoffs $p_{\text{non-sig}} \in \{10^{-6}, 10^{-7}, 10^{-8}, 10^{-10}\}$. We list these measures in Table S3.

#### S5 Application to Mean Corpuscular Haemoglobin PGSs

The following summaries of mean corpuscular haemoglobin (MCH) PGSs obtained from the PGS Catalogue are provided.

- Key information about each PGS (Supplementary Table S4)
- Numbers of matching variants and perturbed variants per choice of $p_{\text{non-sig}}$ (Supplementary Table S5)

We next provide a detailed analysis of the MCH PGSs; we recommend reading the Results section of the Main Text beforehand. As noted in the Main Text, MCH is a biomarker for anaemia. We chose to analyze this phenotype because our C&T lenient PGS reported the lowest total sensitivity score (4.9, out of a maximum possible score of 11) across the 11 performance metrics in our Performance Relative to pPGS and sPGS experiment. *Note: Here and for the remainder of this Supplement, we define* ***sensitivity*** *as the fraction of* 100 *random projections (e.g., pPGSs or sPGSs) for which the original PGS is more performant for a phenotype.* We obtained 15 MCH PGSs from the PGS Catalogue (Xu et al., 2022; Vuckovic et al., 2020; Tanigawa et al., 2022; Privé et al., 2022; Weissbrod et al., 2022), which differ in the number of variants included, study cohorts and training methodology; see Supplementary Table S4 for a summary. For each PGS, we match variants with GWAS $p$-values computed on our cohort, from which we obtain sets of variants to perturb. We use various $p$-value cutoffs ($p_{\text{non-sig}} \in \{10^{-6}, 10^{-7}, 10^{-8}, 10^{-10}\}$) to define the set of perturbed variants. Because each PGS and each choice of cutoff yielded at least 7 variants to perturb (note $2^7 = 128 > 100$ and $2^6 = 64 < 100$), we performed each perturbation scheme $R = 100$ times to evaluate the sensitivity of the original PGS to both pPGSs and sPGSs. We computed 11 performance metrics as listed in Supplementary Table S1, and evaluated them on the test cohort used in our Performance Relative to pPGS and sPGS experiment.

**Sensitivity scores capture distinct properties of a PGS.** We first compare *performances of the original PGSs* on the test cohort, following a typical assessment of a PGS. With the exception of PGS002371, which was trained using BOLT-LMM on Biobank Japan individuals (Weissbrod et al., 2022), all remaining PGSs are considerably performant when assessing using popular metrics — for example, all remaining PGSs report a percentile-average phenotype rank correlation of at least 0.99, a prevalence within Top 1% scoring individuals of at least 0.86, and a rank correlation with phenotype of at least 0.36. Different PGSs also outperform one another depending on the choice of metric. For example, PGS002339 reports the highest Pearson correlation ($r = 0.60$) and Spearman correlation ($\rho = 0.60$) with phenotype and the highest odds ratio for the Top 10% scoring individuals (OR = 12.4), whereas PGS002206 reports the highest cosine similarity with phenotype (CosSim = 0.51) and PGS001989 reports the highest percentile-average phenotype rank correlation ($\rho = 0.999$). We summarize the comparative performances of the PGSs by ranking them for each performance metric and summing the ranks across all 11 metrics (Supplementary Figure S3). Based on this aggregation, which also included our C&T PGSs, the top three PGSs are PGS002339 (1st), PGS001989 (2nd) and PGS000099 (3rd).

Next, we compare *relative performances of the original PGSs.* Recognizing that there is likely overlap of the training cohorts in each separate PGS with our test cohort, which would favourably bias the performance of the original PGS, we adopt a conservative summary of relative performance. Specifically, we score each original PGS by taking the minimum fraction of random projections (pPGSs or sPGSs) beaten by the original PGS across 11 performance metrics, so that a higher score indicates a PGS is uniformly sensitive to perturbation of effects. When comparing these PGS sensitivity scores, we observed differences in the relative performances of the PGSs. Focusing on relative performance against pPGSs with GWAS $p$-value exceeding $10^{-6}$ (i.e., $p_{\text{non-sig}} = 10^{-6}$), only 9 PGSs report a minimum sensitivity of at least 0.75 (Supplementary Figure S4A). In other words, for each remaining PGS, there is at least one performance metric for which the original PGS performed worse than or equal to 25 pPGSs. The 9 PGSs reporting the highest minimum sensitivities all have at least $10^4$ variants in their corresponding polygenic effect vector, with two PGSs containing close to a million variants (PGS002371 and PGS002705, see Supplementary Table S5). Notably, for the top-performing PGS (see previous paragraph), PGS002339, which contains $1.1 \times 10^6$ variants, the original PGS beat only 0.46 of all pPGSs when performance is measured by cosine similarity with phenotype. Three PGSs report consistently strong sensitivity scores: for PGS001219, PGS001989 and PGS002705, regardless of choice of $p_{\text{non-sig}}$ or the type of random projection considered (pPGS or sPGS), the original PGS beat all 100 random projections, uniformly across all performance metrics. Another notable PGS is PGS002460, which is ranked 10th based on aggregating performance of the original PGS across all performance metrics, but which reports high relative performance (i.e., beat all 100 sPGSs) across all choices of $p_{\text{non-sig}}$ (e.g., see Supplementary Figure S4B). These four PGSs contain between $12,000$ and $850,000$ variants, which is fewer than the $1.1 \times 10^6$ variants included in the most high-dimensional PGS, PGS002339. (See Supplementary Table S5.)

**Sensitivity diagnostic curves provide an interpretable evaluation of variant contribution to PGS.** We next investigate the sensitivity of each PGS as a function of the threshold applied, $p_{\text{non-sig}}$. For each PGS, we sum its sensitivities across all 11 performance metrics to obtain an *aggregate sensitivity.* We find that most aggregate sensitivities exhibit an upward trend as $p_{\text{non-sig}}$ was set to increasingly significant, i.e., smaller, values (Supplementary Figure S4C&D). This is true regardless of the type of random projection considered (pPGS or sPGS). One noticeable exception is PGS002656, which displays a curvilinear trend in aggregate sensitivity under sign flipping (see Supplementary Figure S4D). The trend potentially reflects greater uncertainty in the direction of effect reported for variants included in the PGS whose GWAS $p$-values lie between $10^{-10}$ and $10^{-8}$. Regardless, our observations are consistent with the general views that (1) the effects of the more genome-wide significant variants in a trained polygenic score are less likely to be random (specifically, the effects are *not* permutation-invariant); and (2) the effect directions of the more genome-wide significant variants in a trained polygenic score are more robust. Additionally, between sPGSs and pPGSs, we also find that the majority of PGS report higher aggregate sensitivity to pPGSs. A particularly sharp drop in sPGS sensitivity relative to pPGS sensitivity is seen in PGS002371 (average aggregate sensitivity decrement of $-2.5$ across all cutoffs). These observations are consistent with our finding in Supplementary Material S6 below, that PGS performance is generally more sensitive to pPGSs.

In summary, our sensitivity analyses on MCH PGSs demonstrate that they not only capture orthogonal, meaningful properties of a PGS, but also they can be used to learn statistical properties of sets of variants contributing to the PGS.

#### S6 Sign-flipping versus Permutation Perturbation Sensitivities

To investigate whether, across phenotypes, our C&T PGSs were generally more sensitive to one perturbation scheme rather than the other, we compared pairs of sensitivities (which are in one-to-one correspondence with empirical $p$-values defined in Main Text) of each PGS to sPGSs and to pPGSs. While the strength of the relationship depends on the choice of performance metric (of the 11 available), we find that PGSs are in general more sensitive to effect permutation (pPGSs) than to effect sign flipping (sPGSs). For example, when measuring performance using either percentile-prevalence correlation or percentile-average phenotype correlation — and with either Spearman $\rho$ or Pearson $r$ as correlation statistic — the original PGS beat a greater number of pPGSs than sPGSs, regardless of whether the PGS was trained on the lenient ($p_{\text{PGS}} = 10^{-5}$) or stringent ($p_{\text{PGS}} = 10^{-8}$) model, and the $p_{\text{non-sig}}$ choice (all Wilcoxon signed-rank test $p$-values are significant after Bonferroni correction at FWER = 0.05 threshold; see Supplementary Figures S11 and S12). We also observe significantly greater PGS sensitivity to pPGSs than to sPGSs when Spearman $\rho$ between PGS and phenotype is used, in all settings except for lenient PGSs with $p_{\text{non-sig}} = 10^{-8}$. Given that phenotype correlations, rather than percentile-prevalence correlations, of PGSs appear more sensitive to random projections (Table 1 of Main Text), these results suggest that more care is needed in interpreting the *directionality* of single variant effects in a PGS, especially if the underlying PGS training methodology relies heavily on percentile-prevalence metrics.

#### S7 Relationship of PGS Perturbation Sensitivity to Perturbed-Fixed Architecture

We interrogated differences in genetic architecture between the perturbed variants and the fixed variants of our C&T PGSs, by comparing distributions of effects between the two sets of variants at the single phenotype level and across phenotypes. In doing so, we computed 14 quantities measuring the Perturbed-Fixed Architecture of the trait as estimated by the polygenic effect vector, which we list in Supplementary Table S3. Regardless of the choice of PGS (lenient or stringent) or the perturbation cutoff ($\{10^{-6}, 10^{-7}, 10^{-8}, 10^{-10}\}$), we found that most phenotype PGSs have fixed variant effect sizes that stochastically dominate perturbed variant effect sizes (e.g., for lenient PGSs with $p_{\text{non-sig}} = 10^{-8}$ all 103 phenotypes report Wilcoxon rank-sum test $p$-value $< 0.05$). Across phenotypes, PGSs also have significantly larger maximum, mean and median fixed variant effect sizes than perturbed effect sizes: for example, for lenient PGSs with $p_{\text{non-sig}} = 10^{-8}$ the Wilcoxon signed-rank test $p$-values are $8 \times 10^{-19}$ (maximum), $6 \times 10^{-19}$ (mean) and $6 \times 10^{-19}$ (median). We summarize these statistics in a data file available on Github (see Data Availability in Main Text). These findings show that more significant GWAS hits tend to contribute larger effects to a phenotype's polygenic architecture, as estimated by our PGS models.

To investigate if PGS sensitivity (relative performance to pPGS or sPGS) is driven by architectural features of either the set of fixed variants or the set of perturbed variants, we evaluated relationships between sensitivities of 11 performance metrics and the 14 measures of Perturbed-Fixed Architecture. We computed rank correlations between each "performance metric × architectural feature" pair, and summarize these results in Supplementary Figures S13 and S15. We highlight three particular findings. First, we find salient relationships between the sum of effect sizes of perturbed variants and the sensitivity (to either pPGS or sPGS), when performance is measured by Top 10% Odds Ratio and Top 10% Prevalence (regardless of lenient or stringent PGS model, across all $p_{\text{non-sig}}$ choices, rank correlation $p$-values are significant after Bonferroni correction at FWER = 0.05). Second, the number of perturbed variants is also robustly correlated with sensitivity, also using either Top 10% Odds Ratio or Top 10% Prevalence as performance metric (all rank correlation $p$-values are significant at FWER = 0.05). Third, there is no significant relationship between the empirical variance of the effects of perturbed variants and the sensitivity as measured by any PGS performance metric. These findings show that PGS insensitivity to perturbations is not driven by the extent to which the perturbed variant effects are homogeneous, but is instead driven by the number of genome-wide non-significant variants and their cumulative total effect size.

#### S8 Robustness of PGS Sensitivity Results to Random Projection Construction Methodology

As mentioned in Materials and Methods of the Main Text, we evaluated the performance of an original PGS relative to random projections (pPGSs and sPGSs), by splitting the perturbed PGS vector (i.e., after sign-flipping or permutation of effects is performed) into "$S$ignificant" and "$R$andom" parts,

$$\hat{\boldsymbol{y}} = \underbrace{\sum_{\substack{\text{variant } j \\ \text{non-significant} \\ \text{(perturbed)}}} \beta_j \boldsymbol{x}^{(j)}}_{\hat{\boldsymbol{y}}_R} + \underbrace{\sum_{\substack{\text{variant } j \\ \text{significant} \\ \text{(fixed)}}} \beta_j \boldsymbol{x}^{(j)}}_{\hat{\boldsymbol{y}}_S} .$$

We then construct a "reflected" PGS, $\hat{\boldsymbol{y}}_- = \hat{\boldsymbol{y}}_S - \hat{\boldsymbol{y}}_R$. For a given performance metric $f : (\hat{\boldsymbol{y}}, \boldsymbol{y}) \mapsto \mathbb{R}^+$ (e.g., $f$ could compute rank correlation between $\hat{\boldsymbol{y}}$ and $\boldsymbol{y}$), we report the performance of a random projection as $\max\{f^+, f^-\}$, where $f^+ = f(\hat{\boldsymbol{y}}, \boldsymbol{y})$ and $f^- = f(\hat{\boldsymbol{y}}_-, \boldsymbol{y})$. This **Maximum of Two Scores** approach differs from the straightforward evaluation of performance that considers only $f(\hat{\boldsymbol{y}}, \boldsymbol{y})$, which we shall refer to as the **Usual Construction** (of a performance score). We chose this approach because we expect $\hat{\boldsymbol{y}}_R$ to have equal chance of being positively or negatively correlated with the phenotype, owing to the randomness of its constituent effects $\beta_j$.

To ensure robustness of our findings, for the Performance Relative to pPGSs and sPGSs experiment, we repeated our statistical analyses, except with random projection performance evaluated under the Usual Construction. Recall that these analyses comprise:

(i) Identifying significant relationships between PGS overfitting to population structure and PGS relative performance to random projections (cf. Results in Main Text)

(ii) Identifying if PGSs tend to be more sensitive to sign-flipping or permutation perturbations (cf. Results in Main Text and Supplementary Section S6)

(iii) Identifying significant relationships between Perturbed-Fixed Architecture of polygenic effect vector and PGS relative performance to random projections (cf. Results in Main Text and Supplementary Section S7)

We summarize our findings from these analyses in (i) Supplementary Figures S8 and S10, (ii) Supplementary Figures S11 and S12, and (iii) Supplementary Figures S14 and S16. In brief, empirical $p$-values of relative performance tended to be larger, which is expected given that a random projection's performance under Usual Construction is at most as good as that under Maximum of Two Scores. However, the same qualitative findings are still true. This demonstrates that our reported findings are robust to random projection performance scoring approaches. (Data files are available on Github; see Data Availability in Main Text.)

#### S9 Supplementary Tables and Figures

Table S1: Various measures of performance of PGS used in our experiments. Note that $\hat{\boldsymbol{y}}$ (`y_hat`) denotes the PGS vector, while $\boldsymbol{y}$ (`y`) denotes the phenotype vector. A red check (✓) denotes usage in **Performance Inflation by rPGS** and a green check (✓) denotes usage in **Performance Relative to pPGS and sPGS**.

| **Quantity** | **Definition** | **Experiments Included** |
|---|---|---|
| Cosine similarity with Phenotype | $\text{CosineSim}(\hat{\boldsymbol{y}}, \boldsymbol{y}) = \frac{\hat{\boldsymbol{y}}^T \boldsymbol{y}}{\lvert\lvert\hat{\boldsymbol{y}}\rvert\rvert \cdot \lvert\lvert\boldsymbol{y}\rvert\rvert}$ | ✓ (green) |
| Pearson $r$ with Phenotype | $\text{PearsonR}(\hat{\boldsymbol{y}}, \boldsymbol{y}) = \left\lvert \frac{(\hat{\boldsymbol{y}}-\bar{\hat{\boldsymbol{y}}})^T(\boldsymbol{y}-\bar{\boldsymbol{y}})}{\lvert\lvert\hat{\boldsymbol{y}}-\bar{\hat{\boldsymbol{y}}}\rvert\rvert \cdot \lvert\lvert\boldsymbol{y}-\bar{\boldsymbol{y}}\rvert\rvert} \right\rvert$ | ✓ (green) |
| Spearman $\rho$ with Phenotype | $\text{SpearmanRho}(\hat{\boldsymbol{y}}, \boldsymbol{y})$ = `cor(y_hat,y,method="spearman")` | ✓ (green) |
| Incremental $R^2$ | `null.fit=lm(y~PC1+...+PC20+age+sex)`, `prs.fit=lm(y~PC1+...+PC20+age+sex+PGS)`, compute `prs.fit$R2 - null.fit$R2` | ✓ (red) |
| Prevalence at Top 10% Percentile | Obtain cases and controls by median binarization of $\boldsymbol{y}$, $\text{Prev}^{(10)}(\hat{\boldsymbol{y}}, \boldsymbol{y})$ = fraction of cases in top 10% PGS-scoring individuals | ✓ (green) |
| Prevalence at Top 1% Percentile | Obtain cases and controls by median binarization of $\boldsymbol{y}$, $\text{Prev}^{(1)}(\hat{\boldsymbol{y}}, \boldsymbol{y})$ = fraction of cases in top 1% PGS-scoring individuals | ✓ (green) |
| Odds Ratio for Top 10% Percentile | Obtain cases and controls by median binarization of $\boldsymbol{y}$, $\text{Prev}_{(90)}$ = fraction of cases in bottom 90% PGS-scoring individuals, $\text{OR}_{10}(\hat{\boldsymbol{y}}, \boldsymbol{y}) = \frac{\text{Prev}^{(10)}(1-\text{Prev}_{(90)})}{(1-\text{Prev}^{(10)})\text{Prev}_{(90)}}$ | ✓ (green) |
| Odds Ratio for Top 1% Percentile | Obtain cases and controls by median binarization of $\boldsymbol{y}$, $\text{Prev}_{(99)}$ = fraction of cases in bottom 99% PGS-scoring individuals, $\text{OR}_{1}(\hat{\boldsymbol{y}}, \boldsymbol{y}) = \frac{\text{Prev}^{(1)}(1-\text{Prev}_{(99)})}{(1-\text{Prev}^{(1)})\text{Prev}_{(99)}}$ | ✓ (green) |
| Percentile-Prevalence Correlation | Obtain cases and controls by median binarization of $\boldsymbol{y}$, compute prevalence at each PGS-percentile, $r_{PP}(\hat{\boldsymbol{y}}, \boldsymbol{y})$ = Pearson $r$ of prevalence vector and percentile vector | ✓ (red) ✓ (green) |
| Percentile-Prevalence Spearman $\rho$ | Obtain cases and controls by median binarization of $\boldsymbol{y}$, compute prevalence at each PGS-percentile, $\rho_{PP}(\hat{\boldsymbol{y}}, \boldsymbol{y})$ = Spearman $\rho$ of prevalence vector and percentile vector | ✓ (green) |
| Percentile-Average Phenotype Correlation | Compute average phenotype at each PGS-percentile, $r_{PAP}(\hat{\boldsymbol{y}}, \boldsymbol{y})$ = Pearson $r$ of average phenotype vector and percentile vector | ✓ (green) |
| Percentile-Average Phenotype Spearman $\rho$ | Compute average phenotype at each PGS-percentile, $\rho_{PAP}(\hat{\boldsymbol{y}}, \boldsymbol{y})$ = Spearman $\rho$ of average phenotype vector and percentile vector | ✓ (green) |

Table S2: Various measures of stratification by PC used in our experiments. Note that $\boldsymbol{y}$ (`y`) denotes the phenotype vector or the PGS vector, while $\boldsymbol{PC}_k$ (`PCk`) denotes the $k$th principal component vector. A red check (✓) denotes usage in **Performance Inflation by rPGS** and a green check (✓) denotes usage in **Performance Relative to pPGS and sPGS**.

| Quantity | Definition | Experiments Included |
|---|---|---|
| Unsigned cosine similarity with PC1 | $\text{uCosineSim}(\boldsymbol{y}, \boldsymbol{PC}_1) = \left\lvert \frac{\boldsymbol{y}^T \boldsymbol{PC}_1}{\lvert\lvert\boldsymbol{y}\rvert\rvert \cdot \lvert\lvert\boldsymbol{PC}_1\rvert\rvert} \right\rvert$ | ✓ (red) ✓ (green) |
| Unsigned Pearson $r$ with PC1 | $\text{uPearsonR}(\boldsymbol{y}, \boldsymbol{PC}_1) = \left\lvert \frac{(\boldsymbol{y}-\overline{\boldsymbol{y}})^T (\boldsymbol{PC}_1 - \overline{\boldsymbol{PC}_1})}{\lvert\lvert\boldsymbol{y}-\overline{\boldsymbol{y}}\rvert\rvert \cdot \lvert\lvert\boldsymbol{PC}_1 - \overline{\boldsymbol{PC}_1}\rvert\rvert} \right\rvert$ | ✓ (red) ✓ (green) |
| Unsigned Spearman $\rho$ with PC1 | $\text{uSpearmanRho}(\boldsymbol{y}, \boldsymbol{PC}_1)$ = `abs(cor(y,PC1,method="spearman"))` | ✓ (green) |
| Entropy of unsigned cosine similarities with top 40 PCs | $\sum_{k=1}^{40} \left(\frac{r_k}{r_1+\ldots+r_{40}}\right) \log\left(\frac{r_k}{r_1+\ldots+r_{40}}\right)^{-1}$, where $r_k = \text{uCosineSim}(\boldsymbol{y}, \boldsymbol{PC}_k)$ | ✓ (red) ✓ (green) |
| Entropy of unsigned Pearson $r$ with top 40 PCs | $\sum_{k=1}^{40} \left(\frac{r_k}{r_1+\ldots+r_{40}}\right) \log\left(\frac{r_k}{r_1+\ldots+r_{40}}\right)^{-1}$, where $r_k = \text{uPearsonR}(\boldsymbol{y}, \boldsymbol{PC}_k)$ | ✓ (red) ✓ (green) |
| Entropy of unsigned Spearman $\rho$ with top 40 PCs | $\sum_{k=1}^{40} \left(\frac{r_k}{r_1+\ldots+r_{40}}\right) \log\left(\frac{r_k}{r_1+\ldots+r_{40}}\right)^{-1}$, where $r_k = \text{uSpearmanRho}(\boldsymbol{y}, \boldsymbol{PC}_k)$ | ✓ (green) |
| Rank of unsigned cosine similarity with PC1 | $\text{rank}(\text{uCosineSim}(\boldsymbol{y}, \boldsymbol{PC}_1))$ among $\{\text{uCosineSim}(\boldsymbol{y}, \boldsymbol{PC}_k) : k = 1, \ldots, 40\}$ | ✓ (red) ✓ (green) |
| Rank of unsigned Pearson $r$ with PC1 | $\text{rank}(\text{uPearsonR}(\boldsymbol{y}, \boldsymbol{PC}_1))$ among $\{\text{uPearsonR}(\boldsymbol{y}, \boldsymbol{PC}_k) : k = 1, \ldots, 40\}$ | ✓ (red) ✓ (green) |
| Rank of unsigned Spearman $\rho$ with PC1 | $\text{rank}(\text{uSpearmanRho}(\boldsymbol{y}, \boldsymbol{PC}_1))$ among $\{\text{uSpearmanRho}(\boldsymbol{y}, \boldsymbol{PC}_k) : k = 1, \ldots, 40\}$ | ✓ (green) |
| Model $R^2$ with linear model fitted to top 40 PCs | `my.fit=lm(y~PC1+...+PC40)`, compute `my.fit$R2` | ✓ (green) |
| Model adjusted $R^2$ with linear model fitted to top 40 PCs | `my.fit=lm(y~PC1+...+PC40)`, compute `my.fit$adj.R2` | ✓ (green) |
| No. Significant variables in linear model fitted to 40 PCs | `my.fit=lm(y~PC1+...+PC40)`, compute number of significant variables in `my.fit` | ✓ (green) |

Table S3: Various measures of distributional properties of a PGS computed for our experiments. Metrics are computed for each choice of $p_{\text{non-sig}} \in \{10^{-6}, 10^{-7}, 10^{-8}, 10^{-10}\}$. Note that `betas_fixed` and `betas_perturbed` denote the vector of effects of fixed variants and the vector of effects of perturbed variants, respectively.

| **Quantity** | **Definition** |
|---|---|
| Number of Variants | No. variants with nonzero beta in PGS |
| Number of Perturbed Variants | No. variants with $p \geqslant p_{\text{non-sig}}$ |
| Maximum Effect Size of Fixed Variants | Across all variants $j$ with $p < p_{\text{non-sig}}$, maximum $\lvert\beta_j\rvert$ |
| Median Effect Size of Fixed Variants | Across all variants $j$ with $p < p_{\text{non-sig}}$, median $\lvert\beta_j\rvert$ |
| Average Effect Size of Fixed Variants | Across all variants $j$ with $p < p_{\text{non-sig}}$, mean $\lvert\beta_j\rvert$ |
| Variance of Effect Sizes of Fixed Variants | For all variants $j$ with $p < p_{\text{non-sig}}$, variance of $\lvert\beta_j\rvert$'s |
| Sum of Effect Sizes of Fixed Variants | For all variants $j$ with $p < p_{\text{non-sig}}$, sum of $\lvert\beta_j\rvert$'s |
| Maximum Effect Size of Perturbed Variants | Across all variants $j$ with $p \geqslant p_{\text{non-sig}}$, maximum $\lvert\beta_j\rvert$ |
| Median Effect Size of Perturbed Variants | Across all variants $j$ with $p \geqslant p_{\text{non-sig}}$, median $\lvert\beta_j\rvert$ |
| Average Effect Size of Perturbed Variants | Across all variants $j$ with $p \geqslant p_{\text{non-sig}}$, mean $\lvert\beta_j\rvert$ |
| Variance of Effect Sizes of Fixed Variants | For all variants $j$ with $p \geqslant p_{\text{non-sig}}$, variance of $\lvert\beta_j\rvert$'s |
| Sum of Effect Sizes of Fixed Variants | For all variants $j$ with $p \geqslant p_{\text{non-sig}}$, sum of $\lvert\beta_j\rvert$'s |
| Wilcoxon Test $p$-value | Obtain $S_f$ and $S_p$, sets of effect sizes of fixed and perturbed variants `S_f=abs(betas_fixed)` `S_p=abs(betas_perturbed)` `wilcox.test(S_p,S_f,alternative="greater")$p.value` |
| Ratio of Sums of Effect Sizes (Perturbed vs Fixed) | Obtain $S_f$ and $S_p$, sets of effect sizes of fixed and perturbed variants Compute ratio of sum of elements in each set `ratio=sum(S_p)/sum(S_f)` |

Table S4: Summary of key information about each PGS for mean corpuscular haemoglobin (MCH), including the number of variants reported in the original file, the methodology as reported on the PGS Catalogue, and the publication associated with the PGS. The last two rows report our C&T PGSs to allow for comparison.

| **PGS** | **No. Variants Included** | **Methodology** | **Source** |
|---|---|---|---|
| PGS000099 | 27081 | elastic net<br>Bi-allelic variants (MAF $> 0.01$,<br>INFO $> 0.4$, missingness rate $< 0.1$)<br>with LD thinning ($r^2$ threshold $= 0.5$) | Xu et al. (2022) |
| PGS000174 | 628 | sum of effects of<br>conditionally independent variants<br>obtained from GWAS | Vuckovic et al. (2020) |
| PGS001219 | 13003 | snpnet | Tanigawa et al. (2022) |
| PGS001989 | 44174 | penalized regression | Privé et al. (2022) |
| PGS002206 | 504929 | LDPred2-auto<br>(restricted to HapMap 3 variants) | Privé et al. (2022) |
| PGS002339 | 1109310 | BOLT-LMM | Weissbrod et al. (2022) |
| PGS002371 | 920580 | BOLT-LMM-BBJ | Weissbrod et al. (2022) |
| PGS002411 | 22349 | Pruning and Thresholding (P+T)<br>$p < 0.0001$ | Weissbrod et al. (2022) |
| PGS002460 | 44827 | Pruning and Thresholding (P+T)<br>$p < 0.001$ | Weissbrod et al. (2022) |
| PGS002509 | 151362 | Pruning and Thresholding (P+T)<br>$p < 0.01$ | Weissbrod et al. (2022) |
| PGS002558 | 10888 | Pruning and Thresholding (P+T)<br>$p < 10^{-6}$ | Weissbrod et al. (2022) |
| PGS002607 | 8017 | Pruning and Thresholding (P+T)<br>$p < 5 \times 10^{-8}$ | Weissbrod et al. (2022) |
| PGS002656 | 351633 | PolyFun-pred | Weissbrod et al. (2022) |
| PGS002705 | 979778 | SBayesR | Weissbrod et al. (2022) |
| PGS003560 | 638009 | LDPred2-auto<br>`vec_p_init=seq_log(1e-4,0.9,length.out=10),`<br>`burn_in=100, num_iter=500,`<br>`report_step=5` | Ding et al. (2023) |
| lenient | 372 | Clumping and Thresholding<br>$p < 10^{-5}$<br>restrict to 3 variants per locus | This work |
| stringent | 320 | Clumping and Thresholding<br>$p < 10^{-8}$<br>restrict to 3 variants per locus | This work |

Table S5: Numbers of matching variants and perturbed variants per choice of $p_{\text{non-sig}}$. The last two rows report our C&T PGSs to allow for comparison.

| **PGS** | **No. Matching Variants** | **Matching Fraction** | $\boldsymbol{p_{\text{non-sig}} = 10^{-6}}$ | $\boldsymbol{p_{\text{non-sig}} = 10^{-7}}$ | $\boldsymbol{p_{\text{non-sig}} = 10^{-8}}$ | $\boldsymbol{p_{\text{non-sig}} = 10^{-10}}$ |
|---|---|---|---|---|---|---|
| PGS000099 | 26973 | 0.996 | 25333 | 25535 | 25715 | 25964 |
| PGS000174 | 612 | 0.975 | 82 | 130 | 179 | 252 |
| PGS001219 | 12749 | 0.980 | 11534 | 11730 | 11857 | 12063 |
| PGS001989 | 43991 | 0.996 | 40450 | 40982 | 41312 | 41803 |
| PGS002206 | 503696 | 0.998 | 488632 | 490973 | 492546 | 494759 |
| PGS002339 | 1105604 | 0.997 | 1081785 | 1084335 | 1086042 | 1088464 |
| PGS002371 | 917282 | 0.996 | 899346 | 901455 | 902812 | 904822 |
| PGS002411 | 18339 | 0.821 | 10273 | 11347 | 12092 | 13045 |
| PGS002460 | 34464 | 0.769 | 21925 | 23032 | 23785 | 24743 |
| PGS002509 | 101353 | 0.670 | 61782 | 62903 | 63664 | 64624 |
| PGS002558 | 9179 | 0.843 | 3064 | 3935 | 4606 | 5531 |
| PGS002607 | 6819 | 0.851 | 1416 | 2064 | 2614 | 3479 |
| PGS002656 | 312282 | 0.888 | 230280 | 236372 | 239694 | 243585 |
| PGS002705 | 851626 | 0.869 | 842479 | 844176 | 845172 | 846711 |
| PGS003560 | 636463 | 0.998 | 621608 | 623619 | 624958 | 626829 |
| lenient | 372 | 1.00 | 9 | 21 | 30 | - |
| stringent | 320 | 1.00 | - | - | - | 7 |

### Supplementary Figures

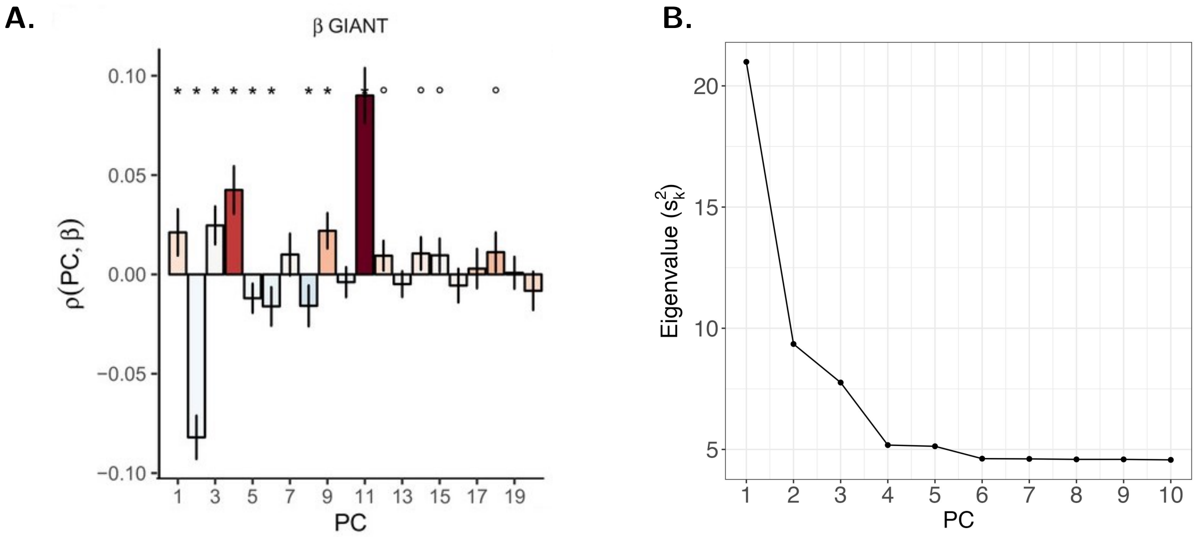

Figure S1: **A.** A plot of Spearman correlations between each PC of the GIANT training cohort and the polygenic effect vector for height. Red bars indicate that the PC has negative Spearman correlation with difference in allele frequencies $\Delta_{\mathrm{AF}}$, between Great British and Toscani Italian populations from the 1000 Genomes Project, computed across each variant contributing to the polygenic effect vector; while blue bars indicate a positive correlation. Colour depth measures the magnitude of the $\Delta_{\mathrm{AF}}$-PC Spearman correlation. Error bars depict 95% bootstrap confidence intervals. Figure modified from Figure 2 of Sohail et al. (2019), in adherence with the Creative Commons Attribution license guidelines. **B.** Empirical eigenvalues computed from running FastPCA on 113,851 UK Biobank samples of UK ancestry and 202,486 SNPs. Figure generated from values reported in Supplementary Table S2 of Galinsky et al. (2016).

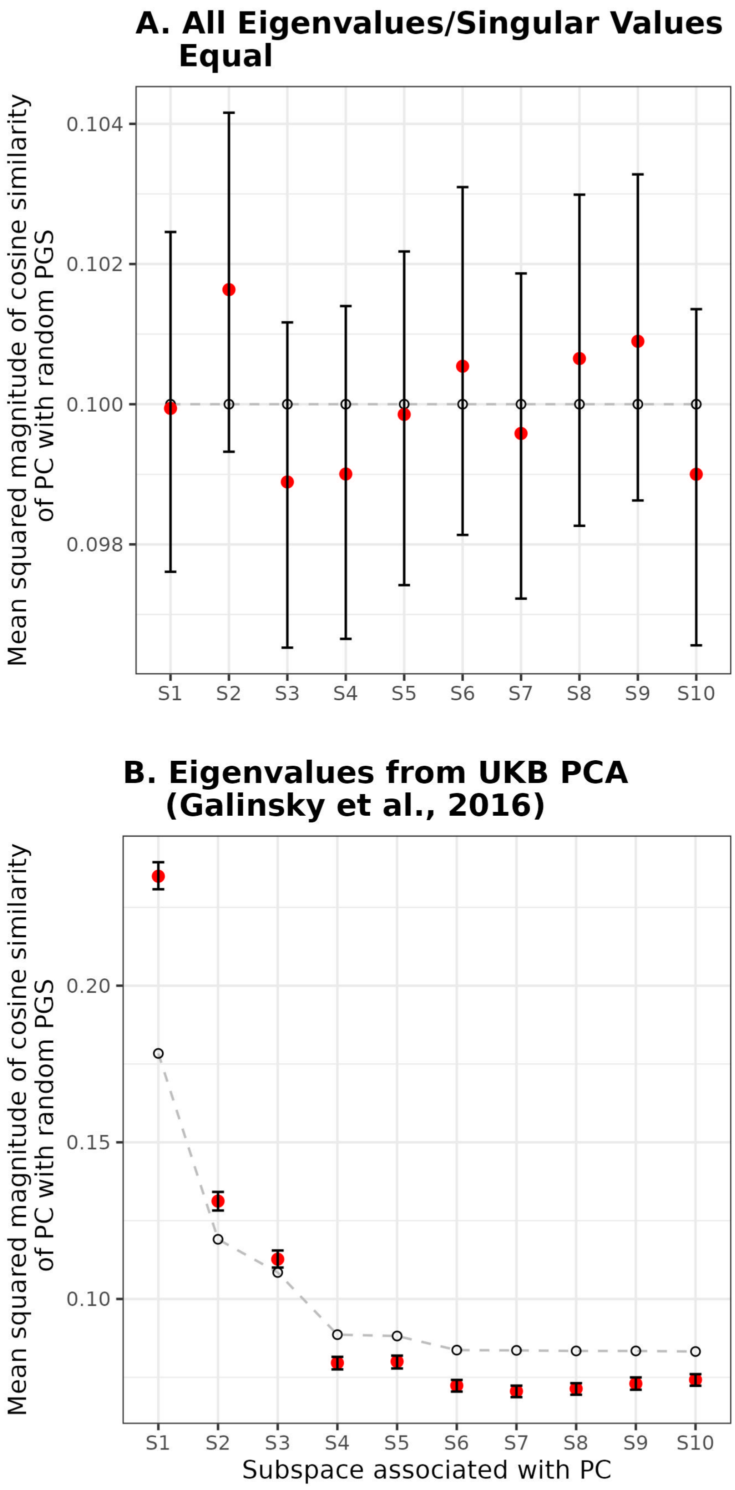

Figure S2: Plot of $\text{Var}(\ell_k)$ against subspace $k$ for $k = 1, \ldots, r$. Here $r = 10$ is assumed for visualization purposes. Analytical predictions ($s_1/(s_1 + \ldots + s_r)$, marked by $\circ$) are plotted against empirical estimates from $10^4$ Monte Carlo draws (marked by $\bullet$, with bootstrap 95% CI). **A.** Case where all singular values $s_k$ are equal to 1. **B.** Case where singular values are set to the squareroot of eigenvalues reported for UKB PCA in Galinsky et al. (2016).

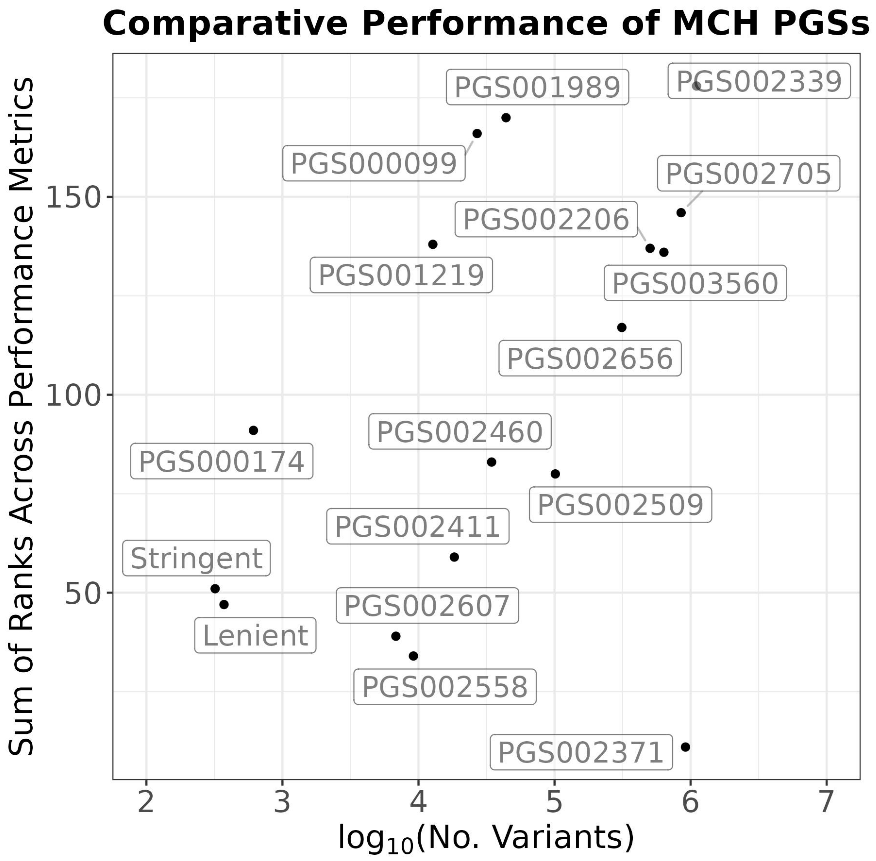

Figure S3: Comparative performance of PGSs for Mean Corpuscular Haemoglobin (MCH), 15 of which are obtained from the PGS Catalogue. The remaining two are the Lenient and Stringent PGSs developed in our present work, and included for reference. Ranks, across 11 performance metrics, of each PGS are tabulated before being summed together to obtain an aggregate rank score that is shown on the $y$-axis (maximum score $= 17 \times 11 = 187$, minimum score $= 1 \times 11 = 11$). Note that a higher rank reflects better comparative performance of a PGS.

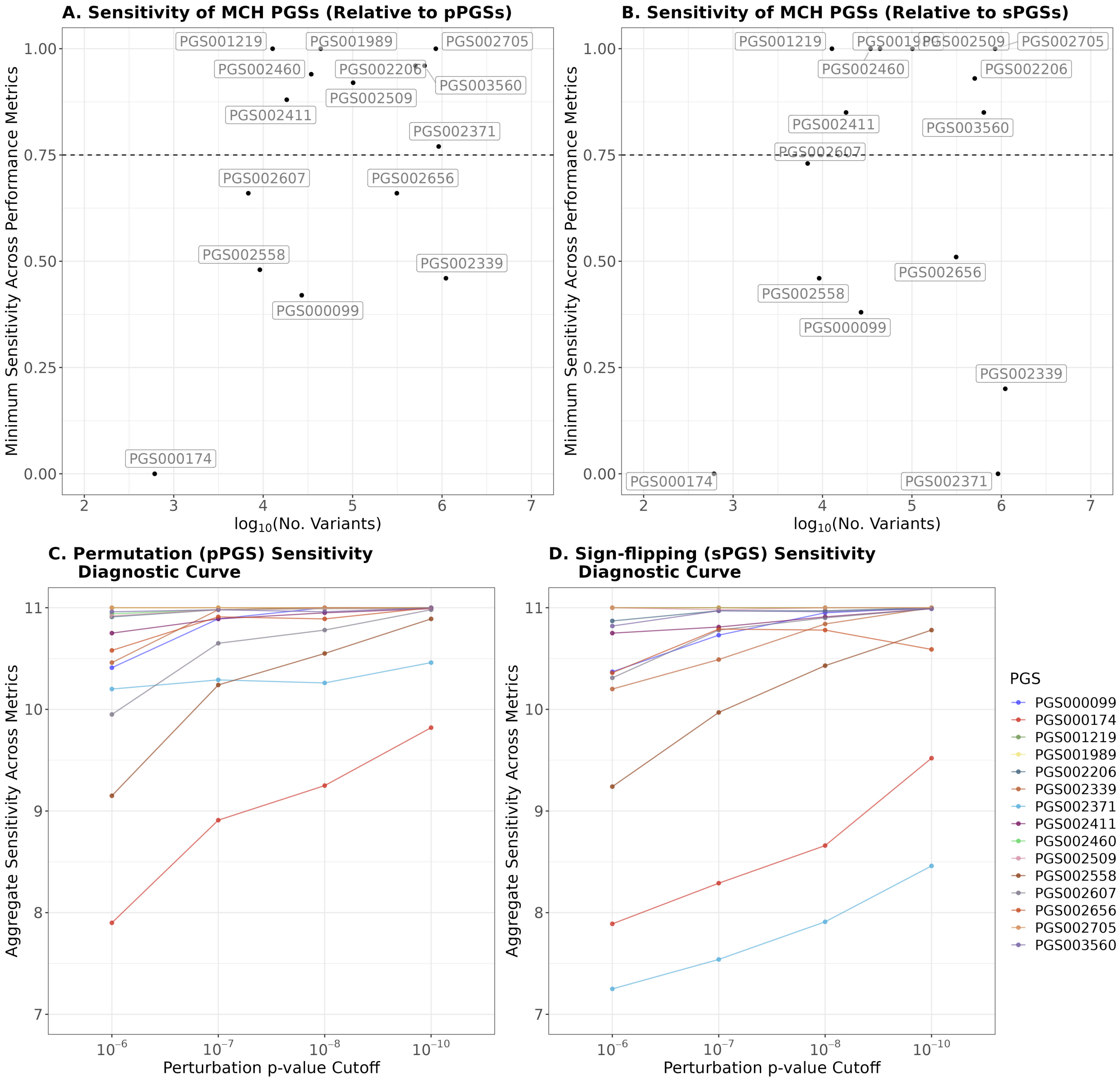

Figure S4: **A.** Minimum sensitivity of MCH PGSs to permutation of non-significant variant effects. In this Figure, non-significant variants are variants with GWAS $p$-value $\geq 10^{-6}$, that is, $p_{\text{non-sig}} = 10^{-6}$ (see Supplementary Material S5 for details). **B.** Minimum sensitivity of MCH PGSs to random sign flipping of non-significant variant effects (see Supplementary Material S5 for details). **C&D.** Permutation (pPGS) sensitivity and sign-flipping (sPGS) sensitivity diagnostic curves for each PGS obtained from the PGS Catalogue. Sensitivities across 11 metrics are normalized and summed for each PGS to obtain its aggregate sensitivity degree (maximum score = 11, minimum score = 0).

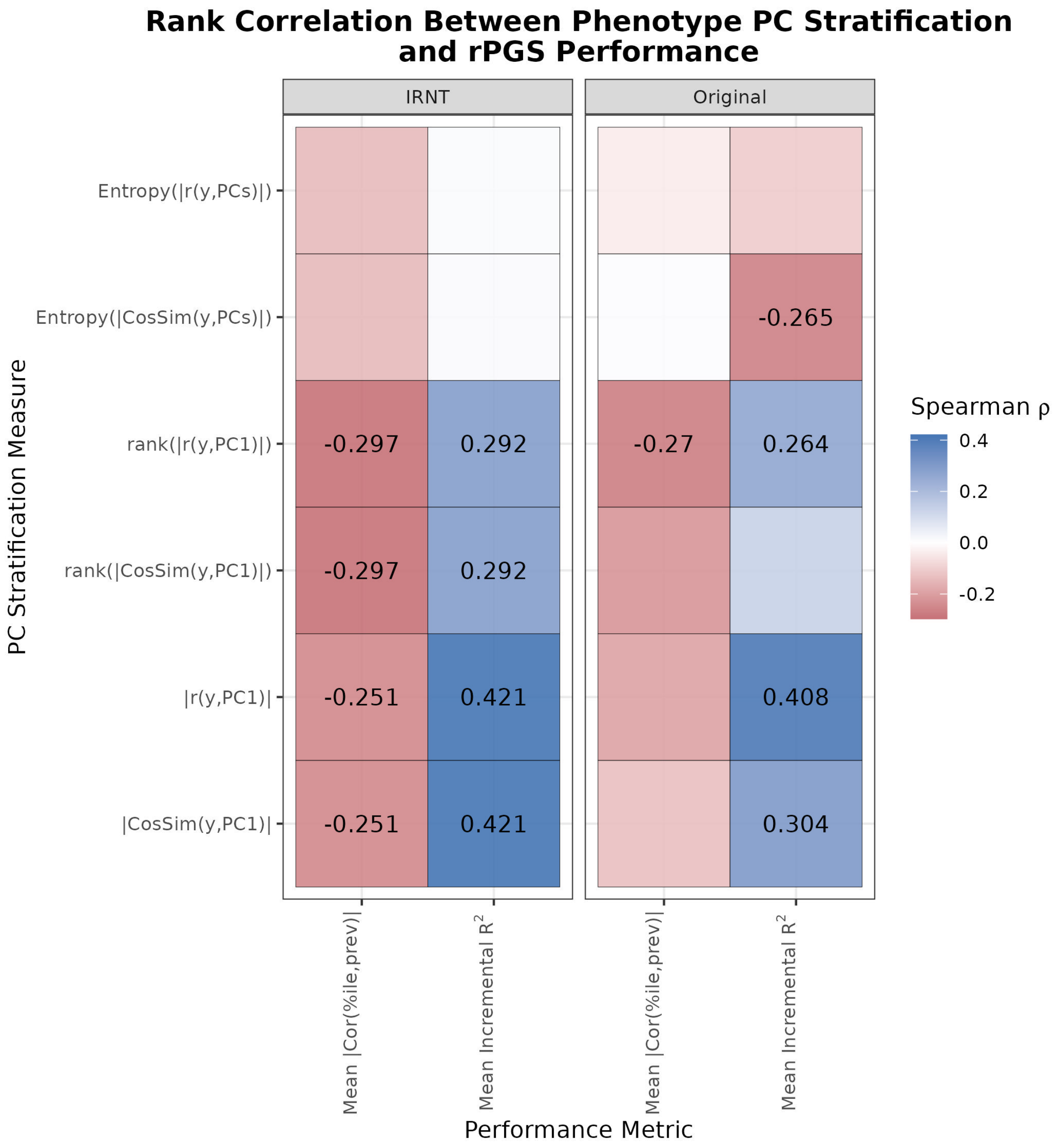

Figure S5: Rank correlations between the performance of 200 rPGSs and PC stratification of a phenotype, across 141 quantitative traits. Performance is measured using either mean magnitude of the correlation between percentile and prevalence based on median-binarizing the phenotype, or the mean incremental $R^2$ against a base model including only the top 20 PCs, age and sex. Only significant (with FWER controlled at 0.05) correlations are shown.

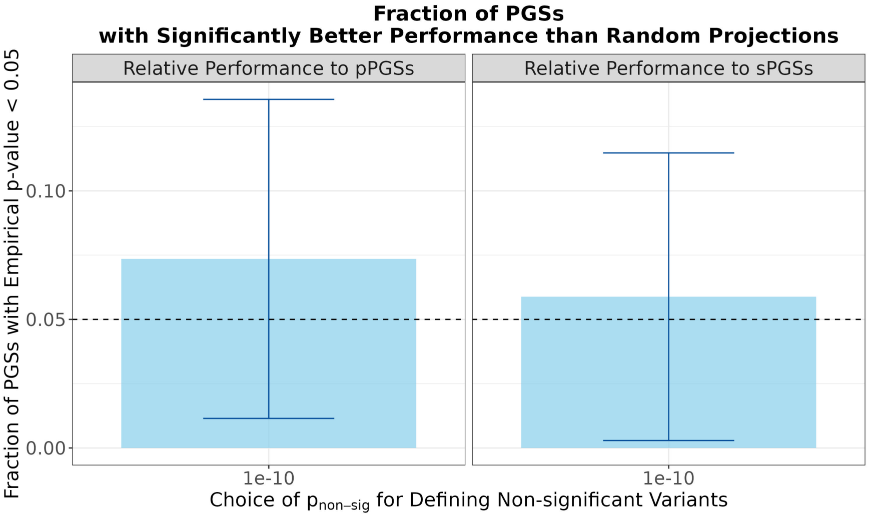

Figure S6: Fraction of PGSs constructed under the stringent model (i.e., including only variants with GWAS $p$-value $< 10^{-8}$) that have relative performance test empirical $p$-value $< 0.05$. Performance is measured by percentile-prevalence rank correlation.

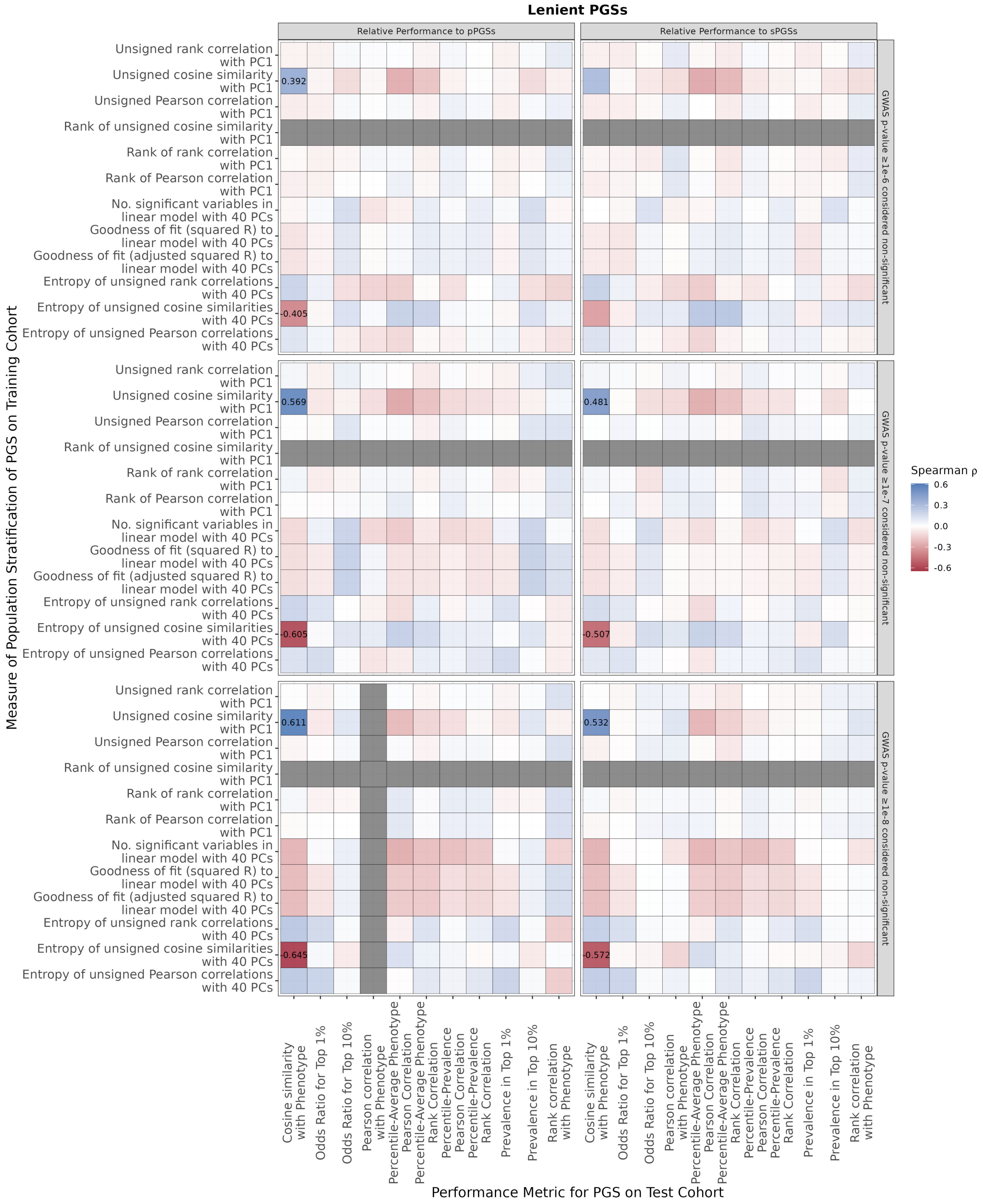

Figure S7: Heatmap of relationships between population stratification of PGS and PGS relative performance empirical $p$-value. For each pair of measure of PGS population stratification and metric of PGS performance (based on which relative performance empirical $p$-values are calculated), the two quantities are computed across all 103 PGSs under the lenient model, before the Spearman correlation is computed. Each cell is coloured by Spearman correlation, with only FWER-controlled significant Spearman correlations shown. FWER control at 0.05 is obtained by applying Bonferroni correction. Gray cells indicate that there is no variation in at least one of the pair of quantities.

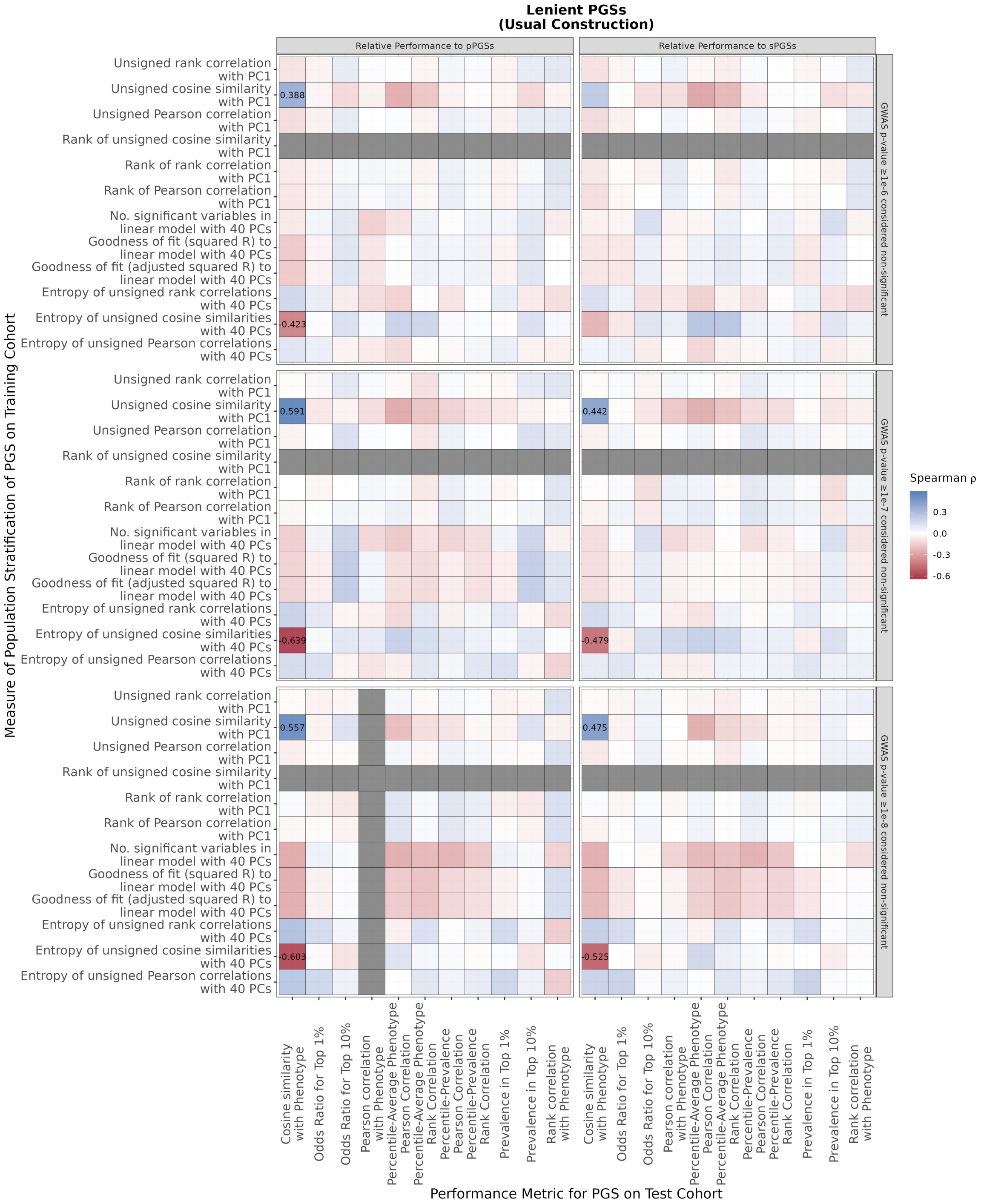

Figure S8: Heatmap of relationships between population stratification of PGS and PGS relative performance empirical $p$-value, under the Usual Construction performance scoring approach (Supplementary Section S8). For each pair of measure of PGS population stratification and metric of PGS performance (based on which relative performance empirical $p$-values are calculated), the two quantities are computed across all 103 PGSs under the lenient model, before the Spearman correlation is computed. Each cell is coloured by Spearman correlation, with only FWER-controlled significant Spearman correlations shown. FWER control at 0.05 is obtained by applying Bonferroni correction. Gray cells indicate that there is no variation in at least one of the pair of quantities.

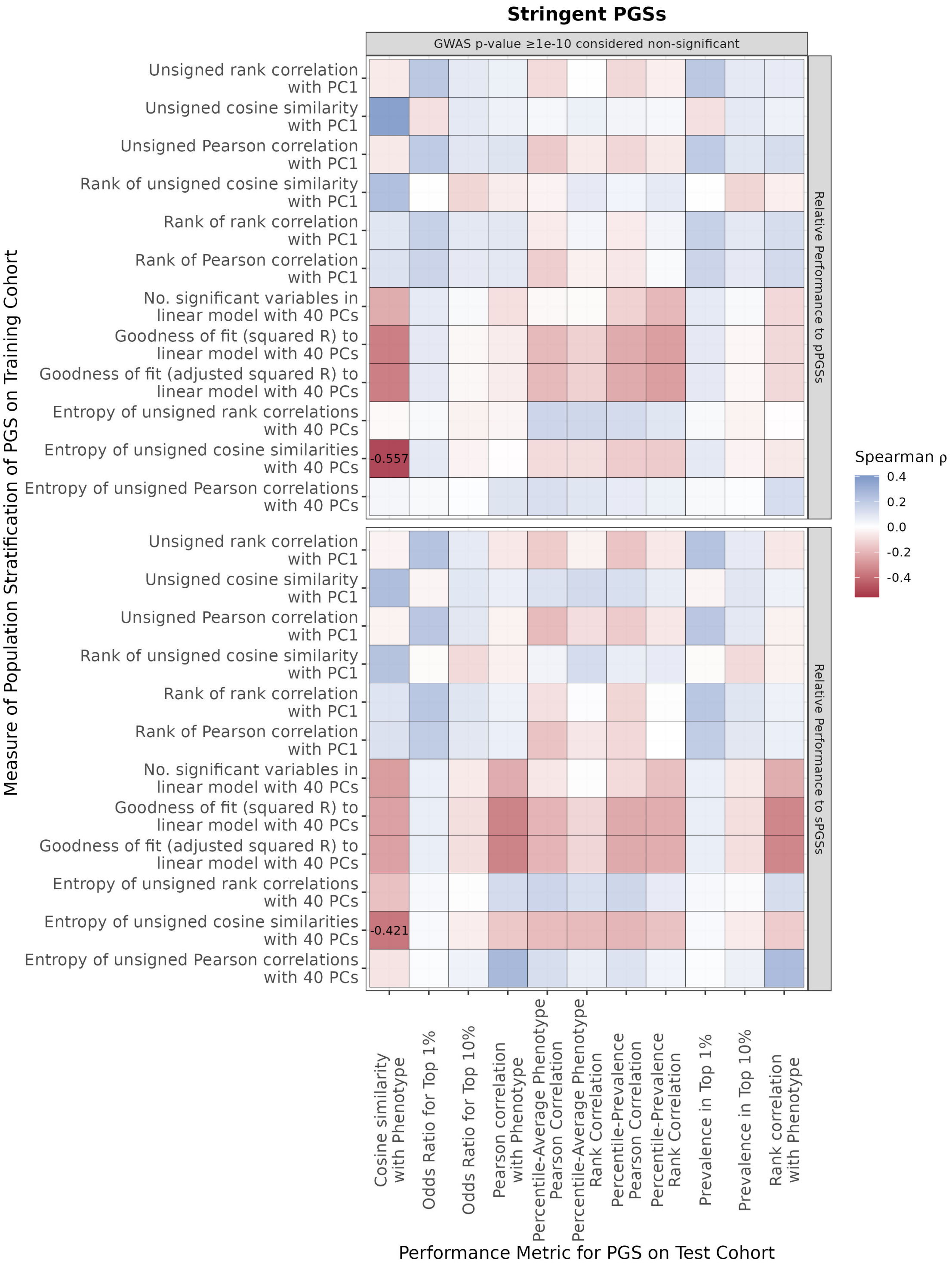

Figure S9: Heatmap of relationships between population stratification of PGS and PGS relative performance empirical $p$-value. For each pair of measure of PGS population stratification and metric of PGS performance (based on which relative performance empirical $p$-values are calculated), the two quantities are computed across all 68 PGSs under the stringent model, before the rank correlation is computed. Each cell is coloured by Spearman correlation, with only FWER-controlled significant Spearman correlations shown. FWER control at 0.05 is obtained by applying Bonferroni correction.

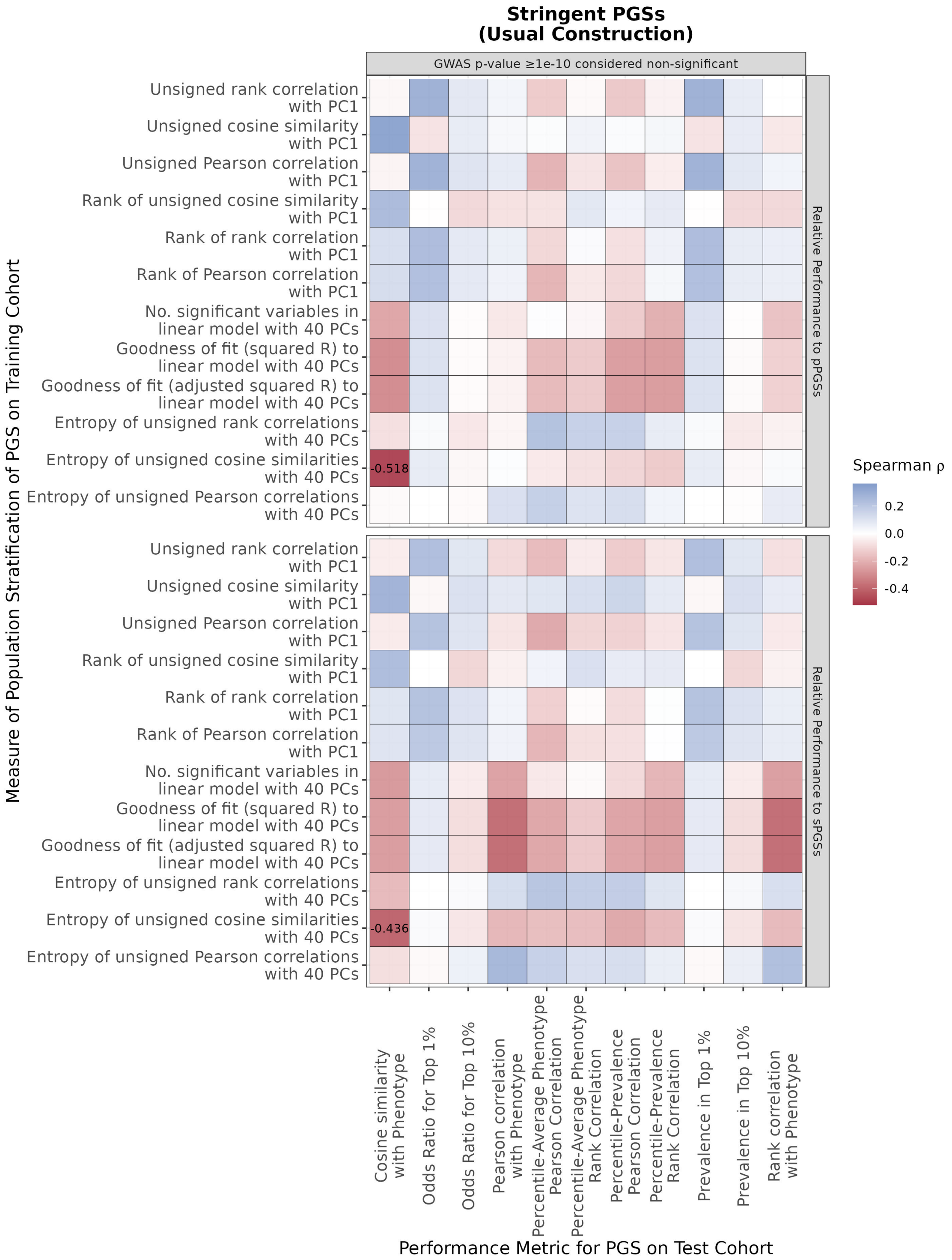

Figure S10: Heatmap of relationships between population stratification of PGS and PGS relative performance empirical $p$-value, under the Usual Construction performance scoring approach (Supplementary Section S8). For each pair of measure of PGS population stratification and metric of PGS performance (based on which relative performance empirical $p$-values are calculated), the two quantities are computed across all 68 PGSs under the stringent model, before the Spearman correlation is computed. Each cell is coloured by Spearman correlation, with only FWER-controlled significant Spearman correlations shown. FWER control at 0.05 is obtained by applying Bonferroni correction.

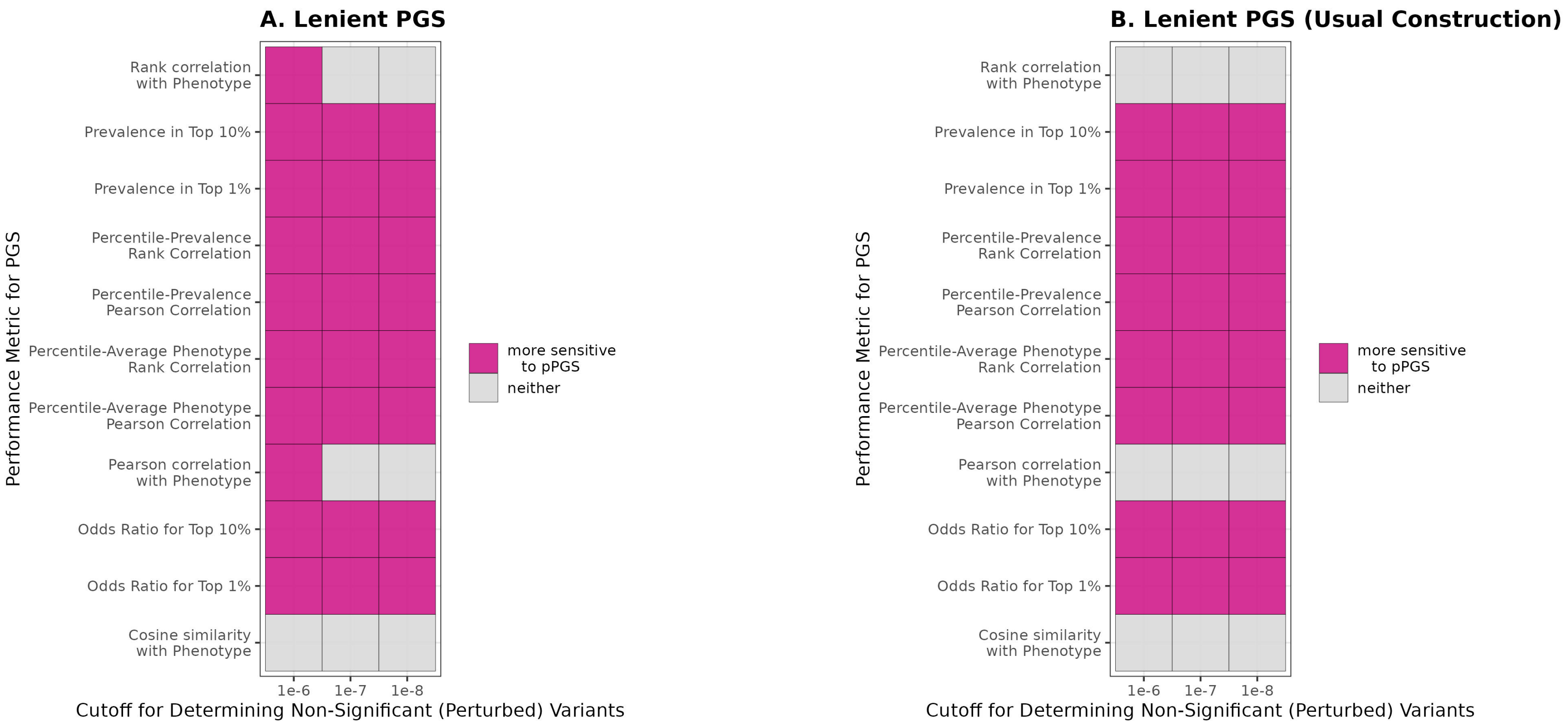

Figure S11: Statistical comparison of sensitivities of original PGSs, under the lenient model, to the two types of random projections, pPGS and sPGS. For each choice of PGS performance metric based on which relative performance of the original PGS was evaluated, a Wilcoxon signed-rank test was performed to determine differential sensitivities. Bonferroni correction was used to control FWER at 0.05 and to identify statistically significant differences. **A.** Performance scoring approach as reported in Main Text. **B.** Performance scoring under Usual Construction (Supplementary Section S8).

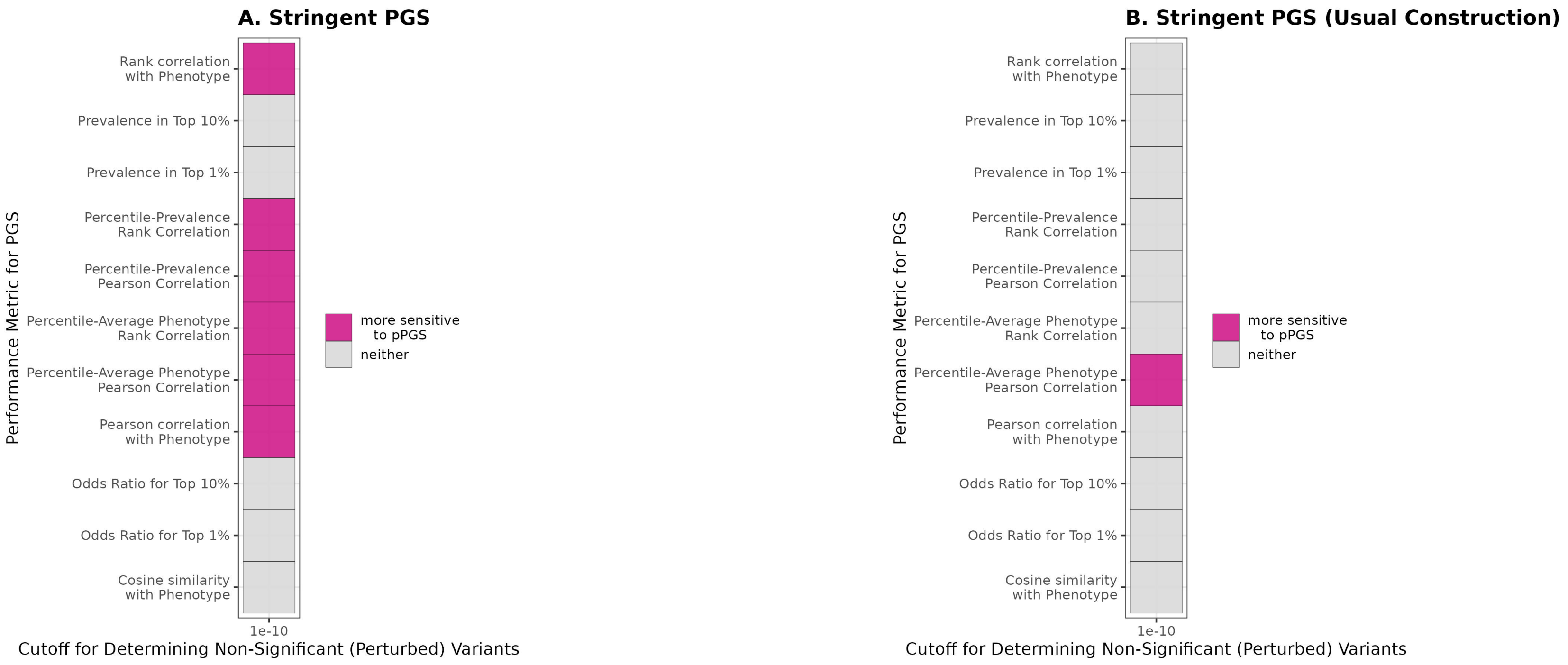

Figure S12: Statistical comparison of sensitivities of original PGSs, under the stringent model, to the two types of random projections, pPGS and sPGS. For each choice of PGS performance metric based on which relative performance of the original PGS was evaluated, a Wilcoxon signed-rank test was performed to determine differential sensitivities. Bonferroni correction was used to control FWER at 0.05 and to identify statistically significant differences. **A.** Performance scoring approach as reported in Main Text. **B.** Performance scoring under Usual Construction (Supplementary Section S8).

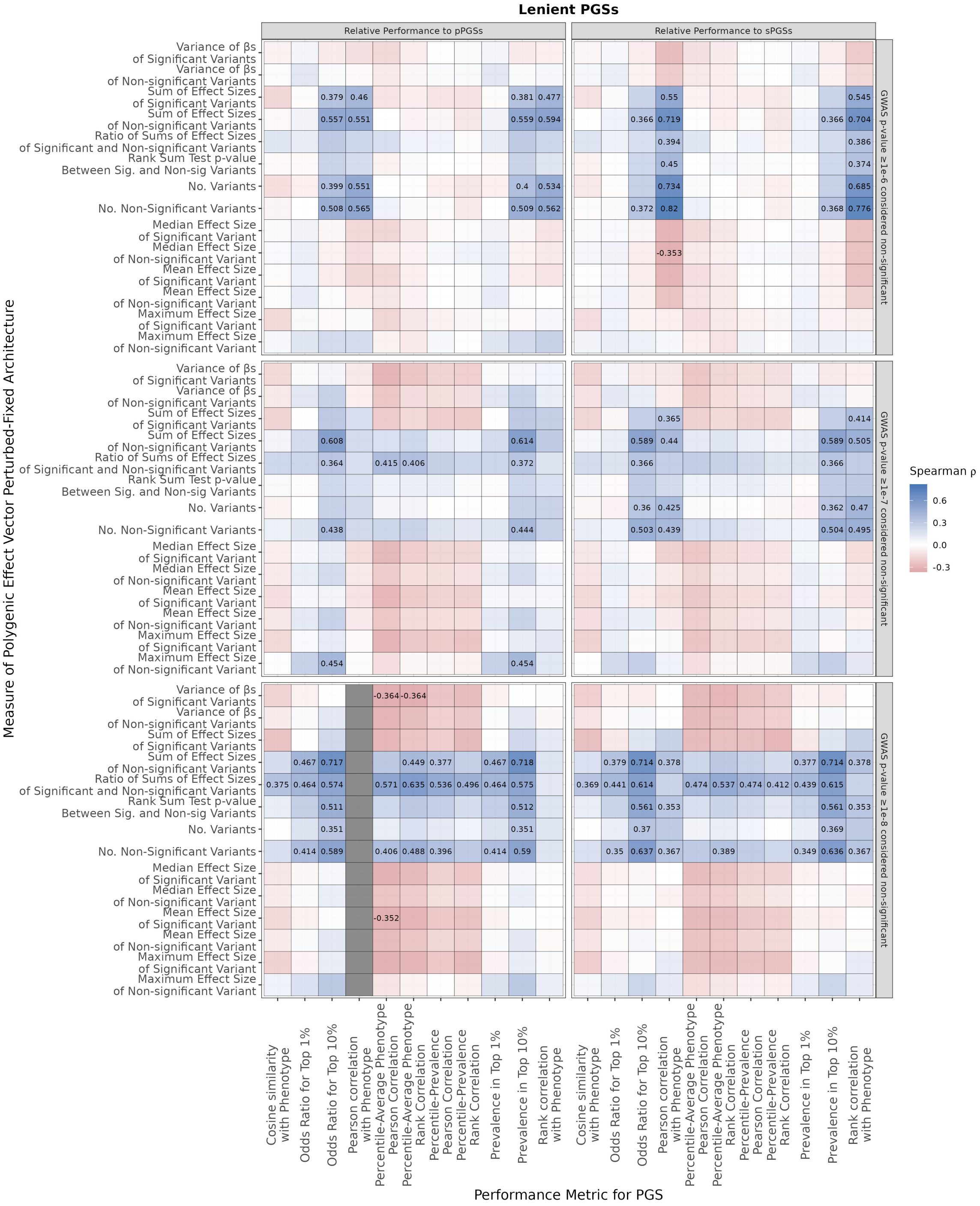

Figure S13: Heatmap of relationships between Perturbed-Fixed Architecture of PGS (see Materials and Methods of Main Text or Supplementary Section S4) and PGS relative performance empirical $p$-value. For each pair of measure of PGS perturbed-fixed architecture and metric of PGS performance (based on which relative performance empirical $p$-values are calculated), the two quantities are computed across all 103 PGSs under the lenient model, before the Spearman correlation is computed. Each cell is coloured by Spearman correlation, with only FWER-controlled significant Spearman correlations shown. FWER control at 0.05 is obtained by applying Bonferroni correction. Gray cells indicate that there is no variation in at least one of the pair of quantities.

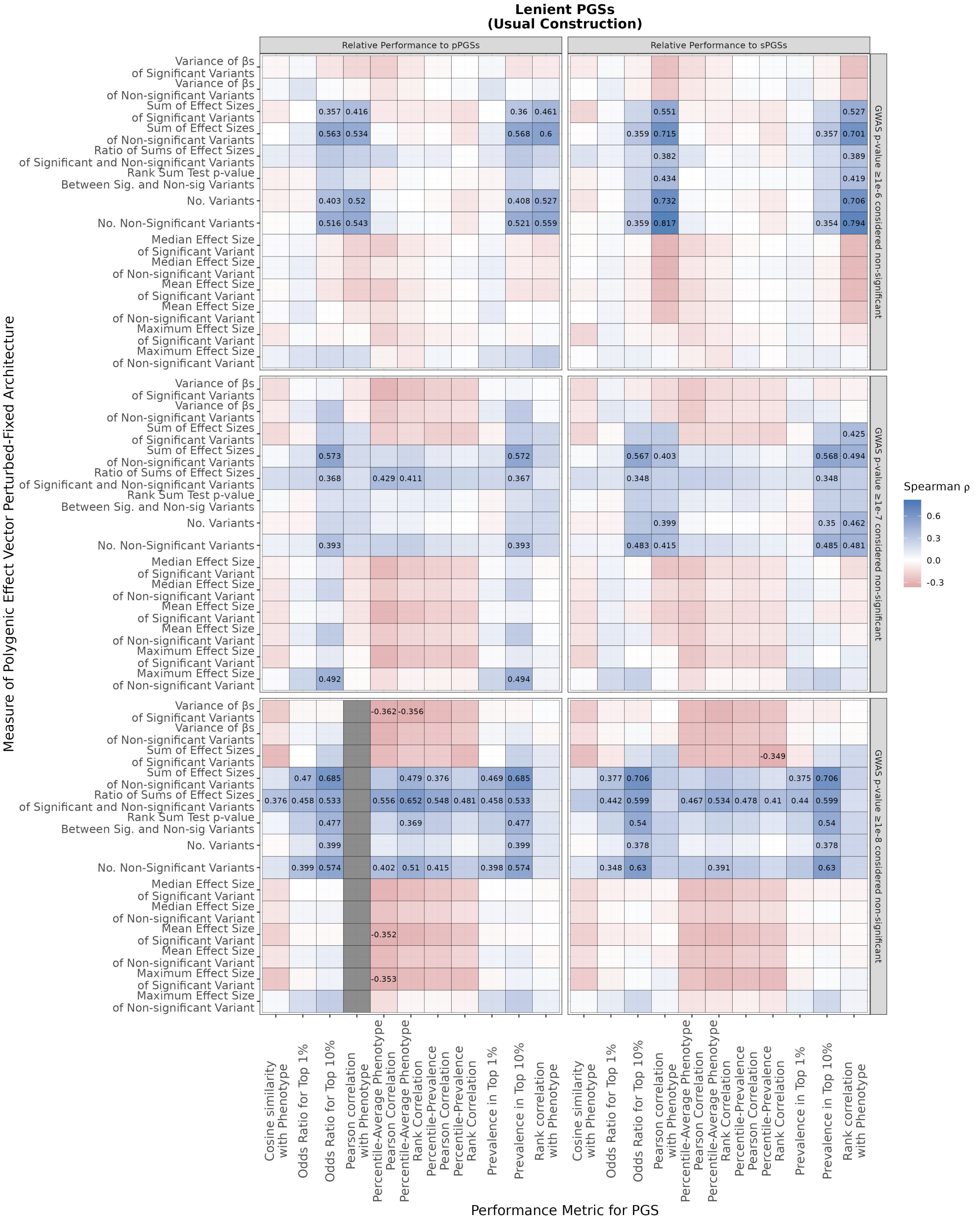

Figure S14: Heatmap of relationships between Perturbed-Fixed Architecture of PGS (see Materials and Methods of Main Text or Supplementary Section S4) and PGS relative performance empirical $p$-value, under the Usual Construction performance scoring approach (Supplementary Section S8). For each pair of measure of PGS perturbed-fixed architecture and metric of PGS performance (based on which relative performance empirical $p$-values are calculated), the two quantities are computed across all 103 PGSs under the lenient model, before the Spearman correlation is computed. Each cell is coloured by Spearman correlation, with only FWER-controlled significant Spearman correlations shown. FWER control at 0.05 is obtained by applying Bonferroni correction. Gray cells indicate that there is no variation in at least one of the pair of quantities.

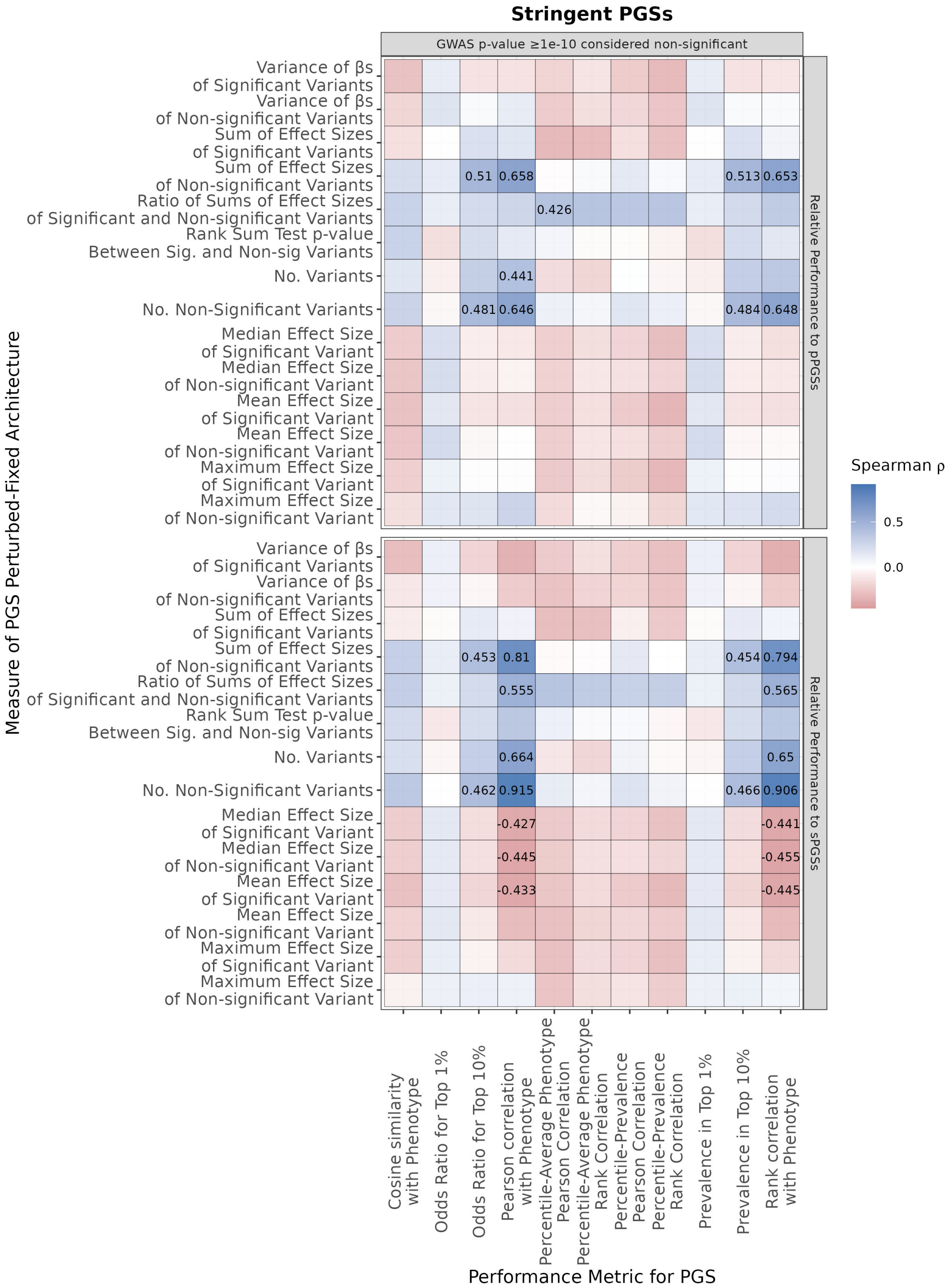

Figure S15: Heatmap of relationships between Perturbed-Fixed Architecture of PGS (see Materials and Methods of Main Text or Supplementary Section S4) and PGS relative performance empirical $p$-value. For each pair of measure of PGS perturbed-fixed architecture and metric of PGS performance (based on which relative performance empirical $p$-values are calculated), the two quantities are computed across all 68 PGSs under the stringent model, before the Spearman correlation is computed. Each cell is coloured by Spearman correlation, with only FWER-controlled significant Spearman correlations shown. FWER control at 0.05 is obtained by applying Bonferroni correction.

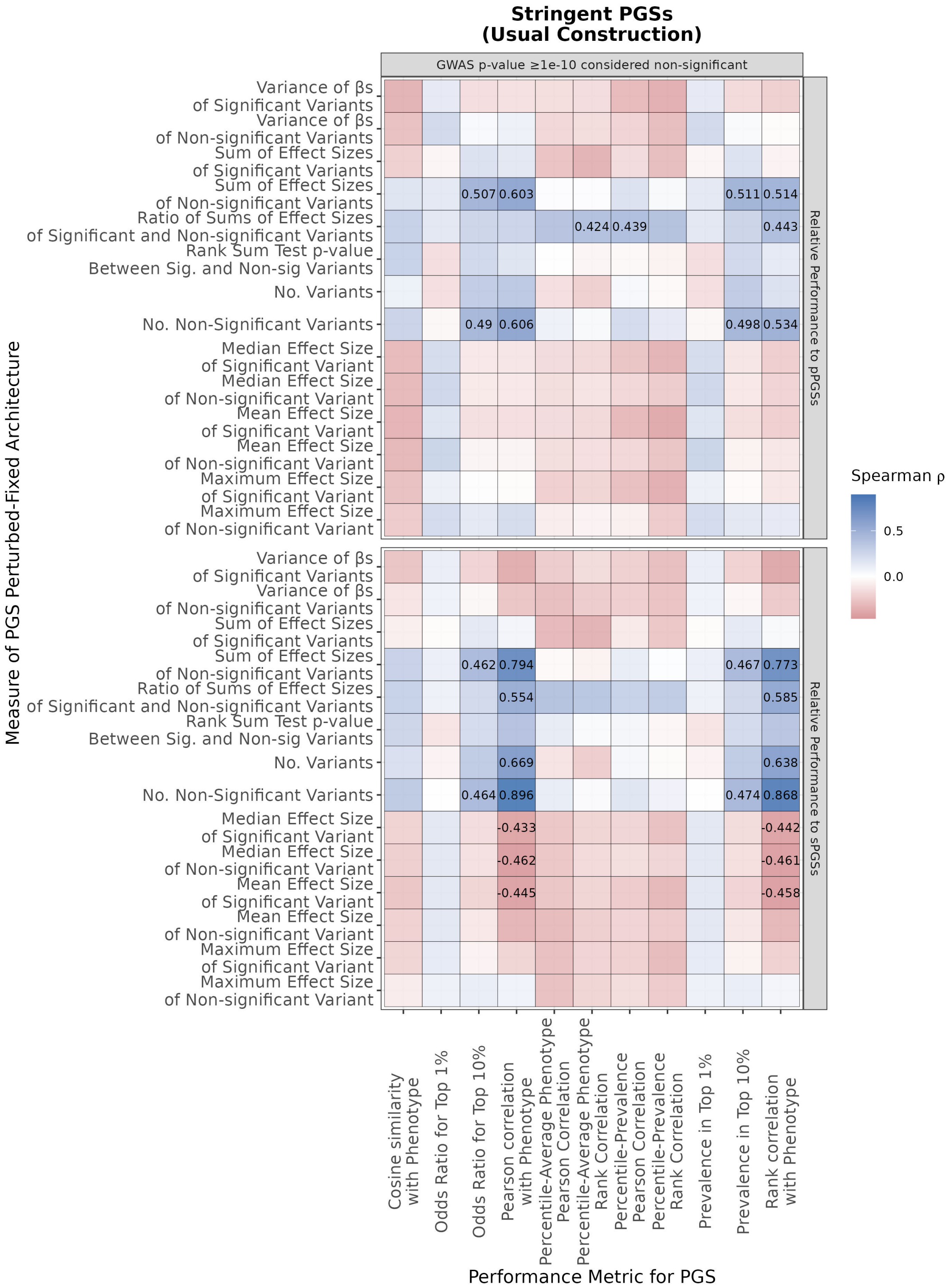

Figure S16: Heatmap of relationships between Perturbed-Fixed Architecture of PGS (see Materials and Methods of Main Text or Supplementary Section S4) and PGS relative performance empirical $p$-value, under the Usual Construction performance scoring approach (Supplementary Section S8). For each pair of measure of PGS perturbed-fixed architecture and metric of PGS performance (based on which relative performance empirical $p$-values are calculated), the two quantities are computed across all 68 PGSs under the stringent model, before the Spearman correlation is computed. Each cell is coloured by Spearman correlation, with only FWER-controlled significant Spearman correlations shown. FWER control at 0.05 is obtained by applying Bonferroni correction.

#### References

Milton Abramowitz, Irene A Stegun, and Robert H Romer. *Handbook of mathematical functions with formulas, graphs, and mathematical tables.* American Association of Physics Teachers, 1988.

Clare Bycroft, Colin Freeman, Desislava Petkova, Gavin Band, Lloyd T Elliott, Kevin Sharp, Allan Motyer, Damjan Vukcevic, Olivier Delaneau, Jared O'Connell, et al. Genome-wide genetic data on 500,000 UK Biobank participants. *BioRxiv*, page 166298, 2017.

Yi Ding, Kangcheng Hou, Ziqi Xu, Aditya Pimplaskar, Ella Petter, Kristin Boulier, Florian Privé, Bjarni J Vilhjálmsson, Loes M Olde Loohuis, and Bogdan Pasaniuc. Polygenic scoring accuracy varies across the genetic ancestry continuum. *Nature*, pages 1–8, 2023.

Kevin J Galinsky, Po-Ru Loh, Swapan Mallick, Nick J Patterson, and Alkes L Price. Population structure of UK Biobank and ancient Eurasians reveals adaptation at genes influencing blood pressure. *The American Journal of Human Genetics*, 99(5):1130–1139, 2016.

Eduardo García-Portugués, Davy Paindaveine, and Thomas Verdebout. On optimal tests for rotational symmetry against new classes of hyperspherical distributions. *Journal of the American Statistical Association*, 115(532):1873–1887, 2020.

Kanti V Mardia and Peter E Jupp. *Directional Statistics*, volume 2. Wiley Online Library, 2000.

Florian Privé, Julyan Arbel, and Bjarni J Vilhjálmsson. LDpred2: better, faster, stronger. *Bioinformatics*, 36(22-23):5424–5431, 2020.

Florian Privé, Hugues Aschard, Shai Carmi, Lasse Folkersen, Clive Hoggart, Paul F O'Reilly, and Bjarni J Vilhjálmsson. Portability of 245 polygenic scores when derived from the UK Biobank and applied to 9 ancestry groups from the same cohort. *The American Journal of Human Genetics*, 109(1):12–23, 2022.

Shaun Purcell, Benjamin Neale, Kathe Todd-Brown, Lori Thomas, Manuel AR Ferreira, David Bender, Julian Maller, Pamela Sklar, Paul IW De Bakker, Mark J Daly, et al. Plink: a tool set for whole-genome association and population-based linkage analyses. *The American Journal of Human Genetics*, 81(3):559–575, 2007.

Mashaal Sohail, Robert M Maier, Andrea Ganna, Alex Bloemendal, Alicia R Martin, Michael C Turchin, Charleston WK Chiang, Joel Hirschhorn, Mark J Daly, Nick Patterson, et al. Polygenic adaptation on height is overestimated due to uncorrected stratification in genome-wide association studies. *eLife*, 8:e39702, 2019.

Yosuke Tanigawa, Junyang Qian, Guhan Venkataraman, Johanne Marie Justesen, Ruilin Li, Robert Tibshirani, Trevor Hastie, and Manuel A Rivas. Significant sparse polygenic risk scores across 813 traits in UK Biobank. *PLoS Genetics*, 18(3):e1010105, 2022.

David E Tyler. Statistical analysis for the angular central Gaussian distribution on the sphere. *Biometrika*, 74(3):579–589, 1987.

Dragana Vuckovic, Erik L Bao, Parsa Akbari, Caleb A Lareau, Abdou Mousas, Tao Jiang, Ming-Huei Chen, Laura M Raffield, Manuel Tardaguila, Jennifer E Huffman, et al. The polygenic and monogenic basis of blood traits and diseases. *Cell*, 182(5):1214–1231, 2020.

Omer Weissbrod, Masahiro Kanai, Huwenbo Shi, Steven Gazal, Wouter J Peyrot, Amit V Khera, Yukinori Okada, Alicia R Martin, Hilary K Finucane, et al. Leveraging fine-mapping and multi-population training data to improve cross-population polygenic risk scores. *Nature Genetics*, 54 (4):450–458, 2022.

Yu Xu, Dragana Vuckovic, Scott C Ritchie, Parsa Akbari, Tao Jiang, Jason Grealey, Adam S Butterworth, Willem H Ouwehand, David J Roberts, Emanuele Di Angelantonio, et al. Machine learning optimized polygenic scores for blood cell traits identify sex-specific trajectories and genetic correlations with disease. *Cell Genomics*, 2(1):100086, 2022.

Jian Yang, S Hong Lee, Michael E Goddard, and Peter M Visscher. GCTA: a tool for genome-wide complex trait analysis. *The American Journal of Human Genetics*, 88(1):76–82, 2011.
